# Glycogen synthase kinase 3 alpha/beta deletion induces precocious growth plate remodeling and cell loss in mice

**DOI:** 10.1101/2020.04.04.025700

**Authors:** Supinder Kour Bali, Dawn Bryce, Carina Prein, James R. Woodgett, Frank Beier

## Abstract

Glycogen synthase kinase (GSK) 3 acts to negatively regulate multiple signaling pathways, including canonical Wnt signaling. The two mammalian GSK3 proteins (alpha and beta) are at least partially redundant. While *Gsk3a* KO mice are viable and display a metabolic phenotype, abnormal neuronal development and accelerated aging, *Gsk3b* KO animals die late in embryogenesis or at birth. Selective *Gsk3b* KO in bone delayed development of some bones, whereas cartilage-specific *Gsk3b* KO mice are normal except for elevated levels of GSK3alpha protein. However, the collective role of these two GSK3 proteins in cartilage was not evaluated. To address this, we generated tamoxifen-inducible, cartilage-specific *Gsk3a/Gsk3b* KO in juvenile mice and investigated their skeletal phenotypes. We found that cartilage-specific *Gsk3a/Gsk3b* deletion in young, skeletally immature mice causes precocious growth plate remodeling, culminating in shorter long bones and hence, growth retardation. These mice exhibit inefficient breathing patterns at later stages and fail to survive. The disrupted growth plates in KO mice showed progressive loss of cellular and proteoglycan components and Sox9 positive cells, with increased staining for osteocalcin and type II collagen. In addition, an increase in osteoclast recruitment and cell apoptosis was observed in growth plates. Surprisingly, changes in articular cartilage of *Gsk3a/Gsk3b* KO mice were mild compared to growth plates, signifying differential regulation of articular cartilage vs growth plate tissues. Taken together, these findings emphasize a crucial role of two GSK3 proteins in skeletal development, in particular in the maintenance and function of growth plates.

**Significance:** Growth plate cartilage dynamics determine the rate of endochondral bone growth and thus, our final height. These processes are disturbed in many genetic and acquired diseases, but the intracellular mechanisms responsible for normal growth plate function, as well as the cessation of growth plate activity in puberty, are poorly understood. Here, we demonstrate that specific removal of both GSK3 genes (*Gsk3a* and *Gsk3b*) in postnatal cartilage of mice leads to a severe reduction of endochondral bone growth, premature remodelling of the growth plate, and early death. In contrast, articular cartilage is only mildly affected by deletion of both genes. These studies identify GSK3 signaling as a key regulator of growth plate dynamics and endochondral bone growth.

## INTRODUCTION

The skeleton, the second largest endocrine organ in mammals and home to various cell types, regulates important physiological processes besides providing structure and support (1). During foetal and neonatal life, skeletal development and growth is significantly driven by the process of endochondral ossification. Formation of the growth plate after secondary ossification contributes towards longitudinal bone growth in postnatal life until the end of puberty (2). While most mammals undergo closure of their growth plates at maturity, mice are well known to maintain their growth plates throughout their lives, although with low activity. Therefore, mice serve as a valuable tool to understand the biology of the growth plate and the pathophysiology of related skeletal disorders, which could be exploited to design treatment modalities for abnormal longitudinal growth and short stature.

Glycogen synthase kinase (GSK) 3 is ubiquitously present in all cell types in mammals and is a known negative regulator of canonical Wnt/beta-catenin signaling as well as many other signaling pathways. GSK3 has been extensively studied in the context of cancer, diabetes, neurodegeneration, inflammation and cardiac hypertrophy (3–5). However, limited studies have addressed the role of GSK3 in skeletal development (6,7) and disease (8). Previous work has suggested an interaction, direct or indirect, of GSK3beta with other important signaling molecules such as Wnt/beta-catenin (9), indian hedgehog (IHH) (10,11), and parathyroid hormone related protein (PTHrP) (12). These molecules are also involved in normal skeletal development and bone growth. Dysregulation of these molecules results in abnormalities of growth plate function, including premature closure of growth plate, hence causing short stature and associated disorders in mice (13–16).

Among two GSK3 proteins (alpha and beta), GSK3beta has been more extensively studied. Global *Gsk3b* KO in mice is embryonically lethal (17), whereas bone-specific *Gsk3b* KO mice show delayed development of some bones (6). Cartilage-specific *Gsk3b* KO mice are normal except for increased expression of GSK3alpha, likely compensating for functional loss of GSK3beta in cartilage (7). *Gsk3a* KO mice are viable but exhibit a metabolic phenotype, accelerated aging and abnormalities in neuronal development (5,18,19). To further elucidate the role of GSK3 in cartilage, tamoxifen inducible cartilage-specific *Gsk3a/Gsk3b* KO mice were generated as an *in vivo* tool to study the effects of deletion of both GSK3 proteins in postnatal skeletal development and growth. Our findings show that the absence of both GSK3 proteins in postnatal cartilage causes premature remodeling of the growth plate, skeletal growth retardation, mineralization of the costo-chondral junction of ribs, and subsequent death. These data demonstrate the crucial role of GSK3 signaling in skeletal development, growth and survival of postnatal skeletally immature young mice.

## MATERIAL AND METHODS

### Generation of inducible cartilage-specific Gsk3a/Gsk3b KO mouse strain

Mice were exposed to a 12-hour light–dark cycle, given tap water, and fed regular chow ad libitum. All the experimental mice were handled in accordance to the guidelines from the Canadian Council on Animal Care, and experiments were approved by the Animal Use Subcommittee at The University of Western Ontario.

Generation of mice homozygous for floxed *Gsk3a* (*Gsk3a*^*fl/fl*^) and floxed *Gsk3b* alleles (*Gsk3b*^*fl/fl*^) respectively, have been described elsewhere (18,19). *Gsk3a*^*fl/fl*^ and *Gsk3b*^*fl/fl*^ mice were crossed with C57BL/6 mice to generate mice heterozygous for both the floxed *Gsk3a and Gsk3b* alleles (*Gsk3a*^*fl/wt*^ *Gsk3b*^*fl/wt*^), which were further bred to homozygosity (*Gsk3a*^*fl/fl*^/*Gsk3b*^*fl/fl*^). *Gsk3a*^*fl/fl*^/*Gsk3b*^*fl/fl*^ mice were crossed with mice expressing tamoxifen-inducible *Cre*^*ERT2*+/-^ recombinase under the control of cartilage-specific mouse *Acan* promoter (*AcanCre*^*ERT2*+/-^) (15) to generate cartilage-specific *Gsk3a*^*fl/fl*^/*Gsk3b*^*fl/fl*^ *AcanCre*^*ERT2*+/-^ mice.

### Validation of genotype

All experimental mice involved in this study were bred in-house. Male *Gsk3a*^*fl/fl*^/*Gsk3b*^*fl/fl*^ *AcanCre*^*ERT2*+/-^ mice were routinely bred with female *Gsk3a*^*fl/fl*^/*Gsk3b*^*fl/fl*^ mice to obtain *Gsk3a*^*fl/fl*^/*Gsk3b*^*fl/fl*^ *AcanCre*^*ERT2*+/-^ and *Gsk3a*^*fl/fl*^/*Gsk3b*^*fl/fl*^ littermates according to Mendelian ratio. All experiments were performed using respective littermate controls to minimize possible variations. Genotype of mice was confirmed by PCR using genomic DNA prepared from biopsy samples of ear tissue of weanlings. Presence of floxed *Gsk3a*, floxed *Gsk3b* and *AcanCre*^*ERT2*+/-^ was detected using following primer sets: 5’- CCCCCACCAAGTGATTTCACTGCTA -3’ (forward) and 5’- CTTGAACCTTTTGTCCTGAAGAACC -3’ (reverse) for *Gsk3a*; 5’- GGGGCAACCTTAATTTCATT -3’ (forward) and 5’- GTGTCTGTATAACTGACTTCCTGTGGC -3’ (reverse) for *Gsk3b*; and 5’- GTTATATTCCGGAGCCCACA -3’ (wildtype), 5’- AAAAGCGACAAGAAGACACCA -3’ (common) and 5’- CTCCAGACTGCCTTGGGAAAA -3’ (mutant) for *AcanCre*^*ERT2*+/-^. PCR program comprised of 1 cycle of denaturation at 95°C for 3 min and 35 cycles of denaturation at 95°C for 15 seconds, primer annealing at 60°C for 30 seconds, and extension at 72°C for 60 seconds, followed by infinite hold at 4°C. Amplified PCR products were analyzed on 2% agarose gel, where *Gsk3a* and *Gsk3b* showed a single band of 510 bp and 584 bp, respectively. *AcanCre*^*ERT2*+/-^ mutant mice showed two bands of 200 bp mutant allele and 299 bp wildtype allele, while wildtype littermates showed one band of the wildtype allele. Cartilage-specific deletion of *Gsk3a* and *Gsk3b* in *Gsk3a*^*fl/fl*^/*Gsk3b*^*fl/fl*^ *AcanCre*^*ERT2*+/-^ and *Gsk3a*^*fl/fl*^/*Gsk3b*^*fl/fl*^ mice induced with tamoxifen or vehicle, and was confirmed by immunohistochemical analysis on frontal sections of mouse knee joint.

### Induction of Gsk3a/Gsk3b KO in mice

*Gsk3a*^*fl/fl*^*/Gsk3b*^*fl/fl*^ *AcanCre*^*ERT2*+/-^ and *Gsk3a*^*fl/fl*^*/Gsk3b*^*fl/fl*^ littermates at P28 were administered with tamoxifen (T5648, Sigma-Aldrich) at a dosage of 3mg/20g body weight per day or vehicle (1:10 dilution of 100% ethanol in corn oil) by gavage, daily for five consecutive days. After 72 hours of tamoxifen wash-out period, tamoxifen and vehicle treated mice were euthanized at P36, P43 and P58 for tissue harvest as shown in Fig. 1A for further analyses.

**Fig. 1.**
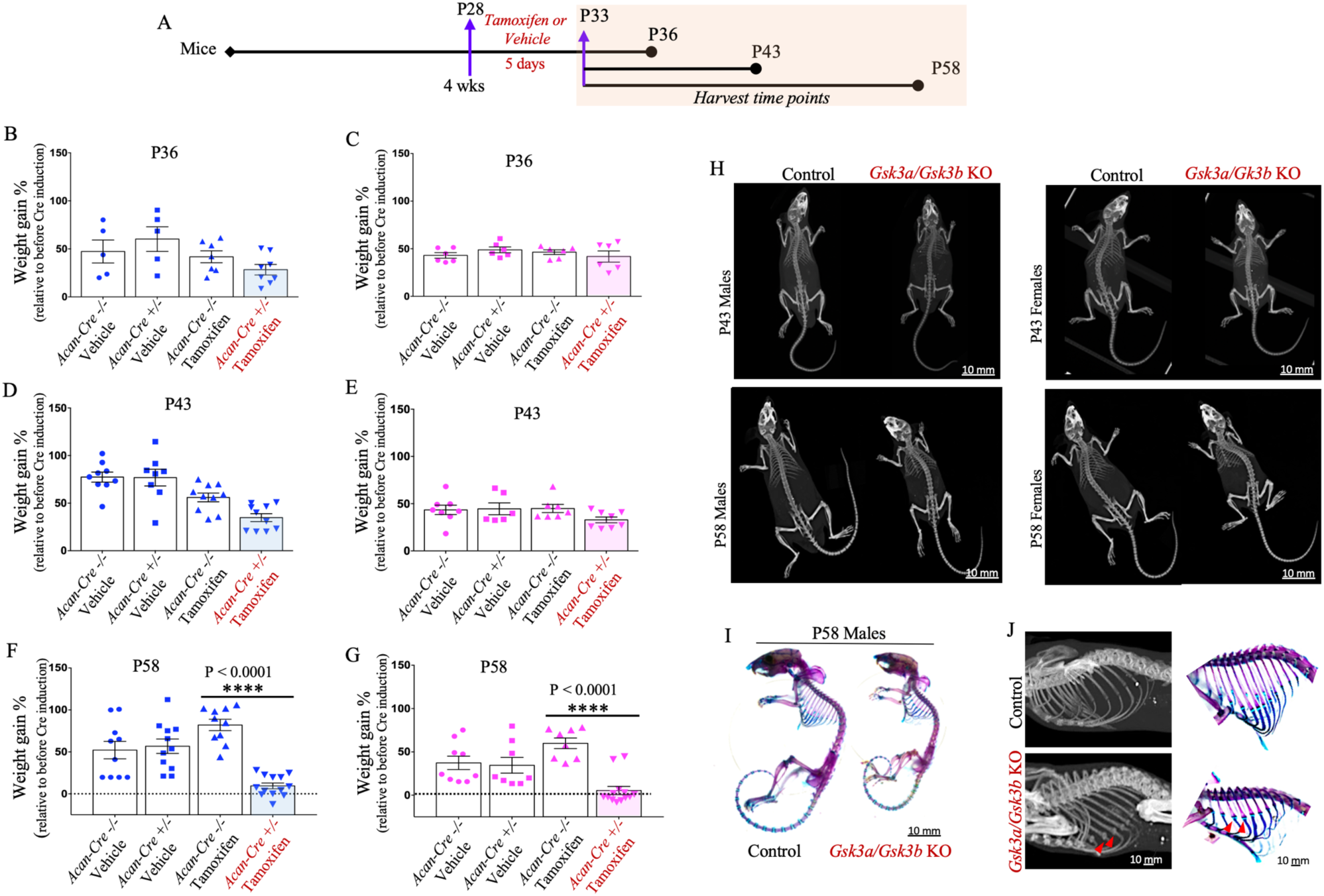
Comparison of growth and skeletal phenotype in young mice after postnatal cartilage-specific *Gsk3a/Gsk3b* KO. Schematic representation of the experimental model used to induce *Gsk3a/Gsk3b* KO in mice and end points for tissue harvest (A). Dot plots with mean show percent weight gain in control and *Gsk3a/Gsk3b* male (B, D, F) and female (C, E, G) KO mice, harvested at P36 (B&C), P43 (D&E) and P58 (F&G). Graphs represent data for individual mouse as well as mean ± SEM from 5-13 mice per experimental group. Statistical significance was calculated using two-way ANOVA with subsequent post hoc Bonferroni test for multiple comparisons. Micro-CT images of male and female *Gsk3a/Gsk3b* KO and control littermates at P43 and P58 (H). Images are representative of 7-13 mice per group. Representative images of six mice per group for Alizarin Red S and Alcian Blue stained mouse skeletons of P58 *Gsk3a/Gsk3b* KO and control littermates (I). Cropped and enlarged region of three dimensional isosurface reconstruction of the mouse ribcage highlighting mineralized costochondral junctions and supporting skeletal staining with red arrows in P58 *Gsk3a/Gsk3b* KO and control littermates (J).

### Micro-computed tomography

Whole body scans for male and female mice were carried out post-partum at P43 and P58. Mice were euthanized and scanned at Robarts Research Institute using a General Electric (GE) Locus Ultra micro-computed tomography (micro-CT) at a resolution of 150μm/voxel. Two-dimensional maximum intensity projection and three-dimensional isosurface images generated using Microview 2.5.0-3943 software (Parallax Innovations, Ilderton, ON, Canada) were used to examine skeletal phenotype and to measure long bone lengths.

### Skeletal staining with Alizarin Red S and Alcian Blue

Euthanized mice at P58 were processed for staining of bone and cartilage components of the whole skeleton as previously described (20,21). Representative images of six independent trials of *Gsk3a*^*fl/fl*^/*Gsk3b*^*fl/fl*^ *AcanCre*^*ERT2*+/-^ and *Gsk3a*^*fl/fl*^*/Gsk3b*^*fl/fl*^ littermate pairs treated with tamoxifen or vehicle are shown.

### Histological analysis

Knee joints were isolated from euthanized mice, fixed in 4% paraformaldehyde, decalcified in EDTA, paraffin embedded, and sectioned at 5μm thickness. Histological examination was performed on 8-10 frontal sections spanning through the entire knee joint using Toluidine Blue stain for general histological evaluation of cartilage and proteoglycan content (22); Tartrate-Resistant Acid Phosphatase (TRAP) assay for detection of osteoclast (23); and Picrosirius Red S for fibrillar organization and collagen content in cartilage and bone (24). Images were captured using a Leica DFC295 digital camera and a Leica DM1000 microscope.

### Immunohistochemical analysis

Frontal sections of paraffinized knees joints were used for immunohistochemical analysis. Sections were deparaffinized and re-hydrated as previously described (24). Primary antibodies used for immunolabelling are as follows: goat polyclonal Sox9 (AF-3075, R&D Systems), rabbit monoclonal GSK3alpha (ab40870, Abcam), rabbit polyclonal GSK3beta (12456S, Cell Signaling Technologies), mouse monoclonal type II collagen (Col2; 10R-C135b, Fitzgerald), rabbit polyclonal osteocalcin (ab93876, Abcam), rabbit polyclonal beta-catenin (bs-1165R, Bioss Antibodies) and rabbit polyclonal GLI1 (PA5-72942, Invitrogen, Thermo Fisher). Frontal sections without primary antibody were used as controls. Sections were incubated with respective HRP-conjugated anti-goat, anti-mouse and anti-rabbit IgG, and labelled proteins were probed with diaminobenzidine substrate (K3468, Dako) to reveal positive staining, followed by counterstaining in Methyl Green.

### Terminal deoxynucleotidyltransferase-mediated dUTP Nick End Labeling analysis

Apoptotic chondrocytes in frontal knee joint sections were assessed for Terminal deoxynucleotidyltransferase-mediated dUTP Nick End Labeling (TUNEL) using DNA Fragmentation Detection Kit (QIA33-1KT, Calbiochem) and following manufacturer’s instructions.

### Digestion of glycosaminoglycans on tissue sections

Paraffinized frontal knee joint sections of P58 wildtype mice were deparaffinized, rehydrated and treated with chondroitinase ABC (C2905, Sigma-Aldrich) for 15 minutes at 37°C, followed by Toluidine Blue staining and immunohistochemistry for Col2 as mentioned in above sections.

### Statistics

Comparison of percent body weight gain and long bone lengths is represented as dot plot for individual mouse per group, with mean ± standard error of the mean (SEM). Statistical significance was calculated using two-way ANOVA with a subsequent post hoc Bonferroni test for multiple comparisons. The significance values are defined as *p 0.05, **p 0.01, ***p 0.001, and ****p 0.0001. All statistical analyses were performed using GraphPad Prism (GraphPad Software Inc. La Jolla, CA, USA) v.6.0. Number of mice used in each experimental group per genotype are indicated in figure legends.

## RESULTS

### Postnatal conditional ablation of *Gsk3a/Gsk3b* in chondrocytes causes growth retardation in young mice

Young, actively growing mice at four weeks of age were used to induce cartilage-specific inactivation of *Gsk3a/Gsk3b* as growth plates at this age are most active and these animals are yet to attain skeletal maturity. Inducible, cartilage-specific *Gsk3a/Gsk3b* KO mice were generated by breeding *AcanCre*^*ERT2*+/-^mice (15) with *Gsk3a*^*fl/fl*^*/Gsk3b*^*fl/fl*^ mice, yielding *Gsk3a*^*fl/fl*^*/Gsk3b*^*fl/fl*^ *AcanCre*^*ERT2*+/-^offspring. These offspring were gavaged with tamoxifen to activate Cre-recombinase in *Gsk3a*^*fl/fl*^*/Gsk3b*^*fl*/fl^ *AcanCre*^*ERT2*+/-^ mice and induce cartilage-specific *Gsk3a/Gsk3b* KO, while Cre-negative *Gsk3a*^*fl/fl*^*/Gsk3b*^*fl/fl*^ littermates were used as controls. A second set of control mice for genotype *Gsk3a*^*fl/fl*^*/Gsk3b*^*fl/fl*^ *AcanCre*^*ERT2*+/-^ and *Gsk3a*^*fl/fl*^*/Gsk3b*^*fl/fl*^ mice treated with vehicle were included throughout the study. Mice of these four groups were evaluated for all assays included in this study and representative figures show articular cartilage and growth plate from the medial compartment for consistency. For clarity, data from vehicle-treated animals are shown in supplementary information.

Four week old mice (P28) of each genotype received tamoxifen or vehicle for five consecutive days by gavage. Treated mice were euthanized at P36, P43 and P58 (Fig. 1A) and analyzed for the effect of postnatal cartilage-specific *Gsk3a/Gsk3b* KO on their growth. Using percent body weight measurements, slightly lower body weight gain was observed at P36 (Fig. 1 B&C) and P43 (Fig. 1 D&E) in KO mice compared to pre-gavage weight (P25), but the effect was not statistically significant. In comparison, KO mice harvested at P58 showed a significant decrease in relative body weight gain in both male (Fig. 1F) and female mice (Fig. 1G). In fact, most KO mice had returned to baseline weights by P58. In addition, the KO animals appeared significantly smaller in size, lethargic, and less mobile, exhibited short-frequent breathing patterns, and did not survive after P59. No such symptoms were visible in P36 and P43 KO mice, or any controls.

### Inducible cartilage-specific *Gsk3a/Gsk3b* KO display altered skeletal phenotypes and compromised longitudinal growth in young mice

Skeletal phenotype and long bone lengths were assessed by micro-CT. Whole body micro-CT scans were performed post-mortem at a resolution of 150μm/voxel, for both male and female mice at P43 and P58 time points. Scans showed smaller skeletons in KO mice compared to control littermates, at both time points, in male as well as female mice (Fig. 1H, S1A). Skeletal phenotype and development were further examined using Alcian Blue and Alizarin Red S stains for cartilage and bone components of the skeleton, respectively. Both male and female KO mice appeared developmentally normal, but smaller in size (Fig. 1I, S1B). Surprisingly, mineralized nodule-like structures were observed at costo-chondral junctions (red arrows) in the ribcage of *Gsk3a/Gsk3b* KO mice using micro-CT scans at both P43 (data not shown) and P58, as well as in skeletal staining at P58 (Fig. 1J, S1C).

We also generated three dimensional isosurface reconstruction using 1200 projections for each mouse scanned at a resolution of 150μm/voxel to measure the length of femur, tibia, and humerus using GE MicroView v2.2 software. In both male (Fig. 2 A&C) and female (Fig. 2 B&D) *Gsk3a/Gsk3b* KO mice, all three long bones were significantly shorter compared to control littermates at P43 (Fig. 2 A&B) and P58 (Fig. 2 C&D). Similar phenotypes were seen in other endochondral bones as well (data not shown).

**Fig. 2.**
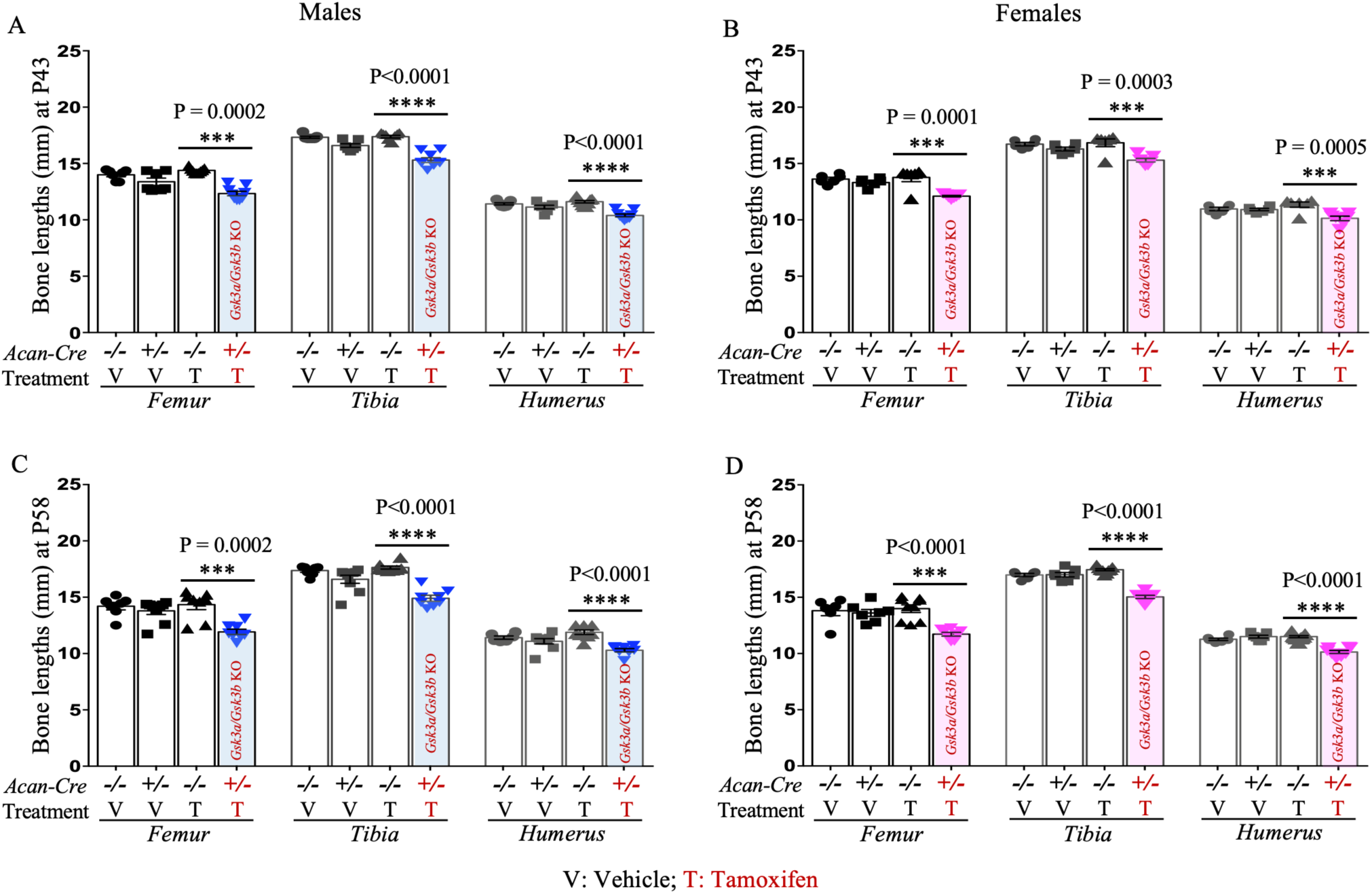
Postnatal cartilage-specific *Gsk3a/Gsk3b* KO mice exhibit shorter long bones. Bone length measurements for femur, tibia and humerus in male (A&C) and female (B&D) *Gsk3a/Gsk3b* KO and control mice at P43 (A&B) and P58 (C&D) were performed using three dimensional isosurface reconstruction of 1200 150u/voxel micro-CT images. Graphs represent measurement for individual mouse as well as mean ± SEM from 5-13 mice per experimental group. Statistical significance was calculated using two-way ANOVA with subsequent post hoc Bonferroni tests for multiple comparisons.

### Cartilage-specific *Gsk3a/Gsk3b* KO mice show loss of structural integrity and precocious remodeling of the growth plate

Our observations demonstrated that *Gsk3a/Gsk3b* KO mice in toto, and their long bones, were smaller in size, but the reason was not known. To address this, histological analyses of the knee joints and cartilage proteoglycan content was performed using Toluidine Blue stained frontal sections in both male and female mice. In *Gsk3a/Gsk3b* KO mice, a progressive loss of structural integrity of the growth plate was observed from P36 to P58 (Fig. 3). Cartilage features in *Gsk3a/Gsk3b* KO mice at P36 included decreased thickness of the growth plate, loss of typical columnar arrangement of chondrocytes and growth plate zonal microarchitecture, drastic decrease in cell number, presence of hypertrophic chondrocytes and formation of chondrocyte cluster-like structures containing four or more cells in the resting zone, and loss of proteoglycan content, as indicated by loss of Toluidine Blue staining. At P43, a significant loss of cellular and proteoglycan content was evident in remnants of growth plate, and these effects were more severe by P58. These changes were consistent in both male (Fig. 3A) and female (Fig. 3B) KO mice across all three time points analyzed, while all control littermates showed normal composition and structural integrity of growth plates (Fig. 3 A&B, S2 A&B). In addition to this growth plate phenotype, a gradual loss of Toluidine Blue stain intensity in articular cartilage was observed in *Gsk3a/Gsk3b* KO mice, in comparison to control mice (Fig. 3 A&B, S2 A&B). However, other than proteoglycan loss, no other degradative symptoms were visible in articular cartilage of the KO mice upon histologic evaluation. Since all of our observations were consistent irrespective of the sex of mice, further results represent data acquired from male mice.

**Fig. 3.**
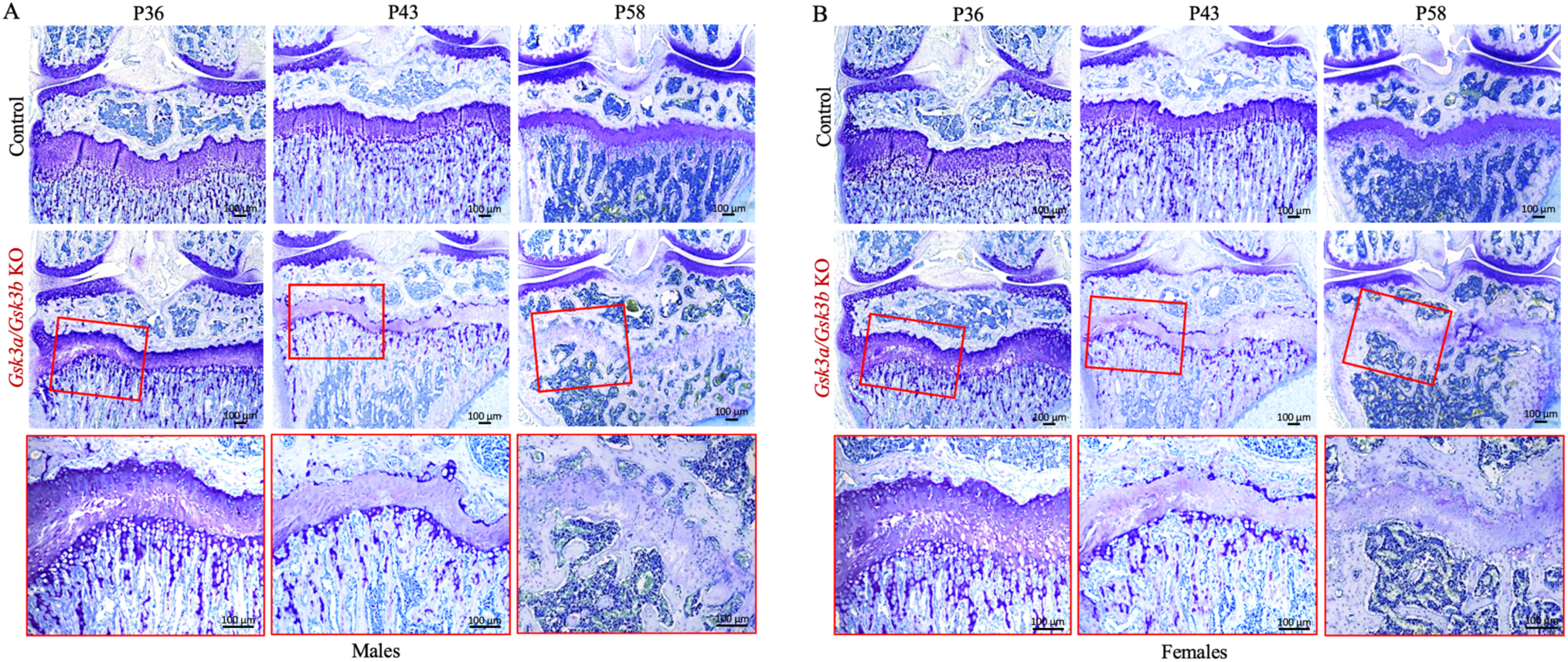
Disorganization of growth plate followed by precocious remodeling in cartilage-specific *Gsk3a/Gsk3b* KO mice. Progressive loss of proteoglycans in cartilage and disorganization of growth plate assessed by Toluidine Blue stained frontal knee joint sections from male (A) and female (B) control and *Gsk3a/Gsk3b* KO mice at P36, P43 and P58 time points. Images are representative of 4-7 mice per group.

### Decreased numbers of Sox9 positive cells in growth plate and articular cartilage of cartilage-specific *Gsk3a/Gsk3b* KO mice

Histological analyses showed loss of cells and proteoglycans in *Gsk3a/Gsk3b* KO cartilage. The cartilaginous nature of growth plate and articular cartilage was further investigated by assessing the presence of Sox9 positive cells. Articular cartilage showed no visible difference in the number of Sox9 positive cells between KO and control littermates at P36, whereas fewer Sox9 positive cells appeared to be present in articular cartilage of *Gsk3a/Gsk3b* KO mice at P43 compared to control littermates. In P58 KO mice, the decrease in the number of Sox9 positive cells was particularly prominent, and positive cells were limited to the superficial zone of articular cartilage compared to control littermates, which had a normal distribution of Sox9 positive cells throughout the articular cartilage (Fig. 4A, S3A). In growth plates of KO mice, Sox9 positive cells were restricted to chondrocytes and chondrocyte-clusters in the resting zone of disorganized growth plate remnants at P36. No Sox9 positive cells were observed in the remnants of the growth plate at P43 and P58 timepoints; however, structures resembling bone marrow cavity-like regions, highlighted by black dotted lines in Fig. 4A, were formed in the region of growth plate remnants at P58, while control littermates did not show these features (Fig. 4A, S3A). Moreover, as mice aged, a gradual decrease in the number of Sox9 positive cells was observed in control mice from P36 through P58 in articular cartilage as well as in the growth plate (Fig. 4A).

**Fig. 4.**
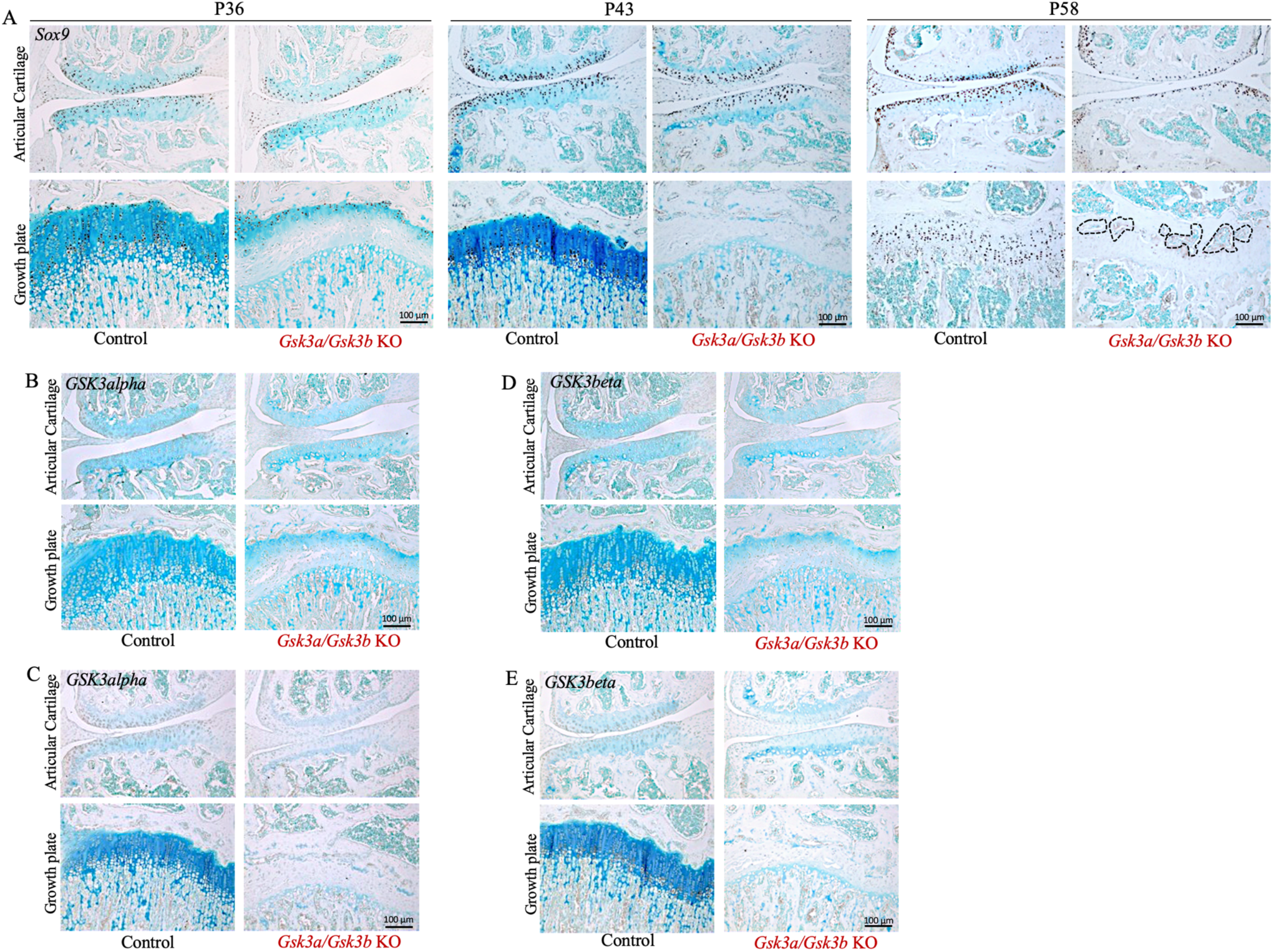
Localized expression of Sox9, GSK3alpha and GSK3beta in cartilage of *Gsk3a/Gsk3b* KO mice. Presence of Sox9 positive cells in *Gsk3a/Gsk3b* KO and control mice at P36, P43 and P58 was assessed by immunohistochemistry on frontal knee joint sections (A). Bone marrow cavity-like regions are highlighted in black dotted line. Localized expression and loss of GSK3alpha (B&C) and GSK3beta (D&E) in *Gsk3a/Gsk3b* KO mice was evaluated by IHC at P36 (B&D) and P43 (C&E). Images are representative of 4-7 mice per group for Sox9, and 3-4 mice per group were used for GSK3alpha and GSK3beta at both time points.

Since the effect of *Gsk3a/Gsk3b* KO was so rapid and drastic, the expression profile of GSK3alpha and GSK3beta proteins in growth plate and articular cartilage was examined in frontal knee joint sections by IHC. At both P36 (Fig. 4 B&C, S3 B&C) and P43 (Fig. 4 D&E, S3 D&E) time points, control mice exhibited expression of both GSK3alpha (Fig. 4 B&D, S3 B&D) and GSK3beta (Fig. 4 C&E, S3 C&E) from the superficial to the middle zones of articular cartilage. While articular cartilage in *Gsk3a/Gsk3b* KO mice showed absence of GSK3 proteins in most areas, faint expression of both GSK3alpha (Fig. 4 B&D) and GSK3beta (Fig. 4 C&E) was observed in a few chondrocytes in the top layer of the superficial zone. While not directly measured, this may be due to the lack of expression or activity of the Cre transgene in these cells. In growth plates of control mice at P36 (Fig. 4 B&C, S3 B&C) and P43 (Fig. 4 D&E, S3 D&E), GSK3alpha was expressed in proliferative to pre-hypertrophic chondrocytes (Fig. 4 B&D, S3 B&D), and GSK3beta was expressed in hypertrophic chondrocytes (Fig. 4 C&E, S3 C&E). No expression of either GSK3alpha (Fig. 4 B&D) or GSK3beta (Fig. 4 C&E) was observed in growth plates of KO mice. This confirmed that gene deletion was effectively carried out by tamoxifen-activated *Acan*-driven Cre recombinase in cartilage tissue of *Gsk3a/Gsk3b* KO mice.

### Accelerated remodeling of growth plate in young mice upon cartilage-specific *Gsk3a/Gsk3b* KO

We next asked what tissue replaced the remodeling growth plate cartilage in *Gsk3a/Gsk3b* KO mice and examined osteogenic differentiation using osteocalcin as a marker for bone formation. Osteocalcin is produced by mature osteoblasts, and subchondral bone matrix showed presence of osteocalcin (Fig. 5A, S4A). Surprisingly, the osteocalcin antibody used showed cellular expression in pre-hypertrophic and hypertrophic chondrocytes of control mice as well, however cartilaginous matrix in both growth plate and articular cartilage did not stain positive for osteocalcin in control mice. In *Gsk3a/Gsk3b* KO mice at P36, the cells present in chondrocyte cluster-like structures stained positively for osteocalcin with no matrix staining in growth plate remnants. In P43 KO animals, no cellular staining for osteocalcin was observed in growth plate remnants due to loss of cellular component, however the intensity of matrix staining appeared increased compared to P36. The remnants of the growth plate showed even more intense osteocalcin staining in P58 *Gsk3a/Gsk3b* KO mice compared to earlier ages, but less intense than mineralized bone (Fig. 5A). These data are consistent with replacement of growth plate tissue by a bone-like matrix. Contrary to the growth plate, articular cartilage matrix in *Gsk3a/Gsk3b* KO mice and respective controls did not show presence of osteocalcin (Fig. 5A, S4A). This implied differential regulation of articular and growth plate chondrocytes.

**Fig. 5.**
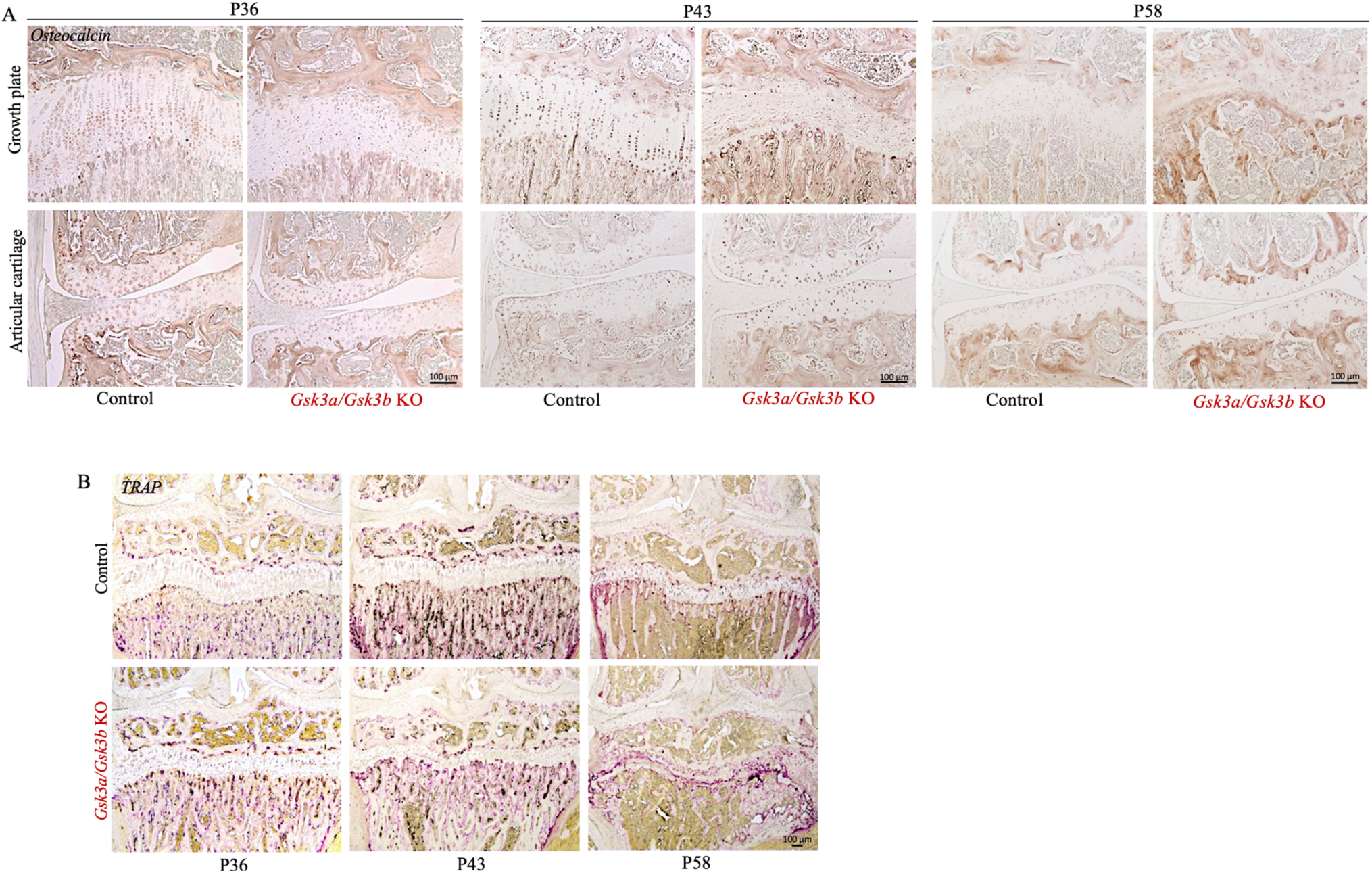
Accelerated remodeling of growth plate in cartilage-specific *Gsk3a/Gsk3b* KO mice. Growth plate remnants in *Gsk3a/Gsk3b* KO mice were examined for the presence of osteocalcin in control and KO mice at P36, P43 and P58 time points (A). Growth plate remodeling was determined by staining for TRAP positive osteoclasts at all three time points (B). Images are representative of 4-7 mice per group for osteocalcin and 3-4 mice per group for TRAP at each time point, respectively.

Growth plate remodeling of *Gsk3a/Gsk3b* KO mice was further assessed by tracing the presence of bone resorbing osteoclasts through tartrate-resistant acid phosphatase (TRAP) staining. Using TRAP assays on frontal knee joint sections, osteoclasts were observed on mineralized bone surfaces in subchondral bone and primary spongiosa under the growth plate in control mice at all three time points (Fig. 5B, S4B). *Gsk3a/Gsk3b* KO mice at P36 showed increased pink staining, indicating more osteoclasts in KO mice compared to control littermates. At P43, *Gsk3a/Gsk3b* KO mice displayed thicker trabeculae and more intense pink osteoclast staining than control littermates as well as P36 KO mice, suggesting increased osteoclast recruitment in KO mice. In P58 *Gsk3a/Gsk3b* KO mice, enhanced osteoclast staining was observed in the remnants of the growth plate compared to control littermates as well as younger KO mice, suggesting increased recruitment of osteoclasts for accelerated resorption of bone-like matrix in the remnants of growth plate. Trabecular microarchitecture in primary spongiosa in P58 *Gsk3a/Gsk3b* KO mice was also drastically resorbed (Fig. 5B, S4B).

### Genetic ablation of *Gsk3a/Gsk3b* in cartilage causes chondrocyte apoptosis and compromises composition of growth plate and articular cartilage

Histological analyses of frontal knee joint sections had shown loss of cells and proteoglycan content in *Gsk3a/Gsk3b* KO cartilage. Cell loss in cartilage can occur because of cell death by apoptosis. TUNEL assays were performed on knee joint sections of P36 mice. No apoptotic cells were seen in articular cartilage of *Gsk3a/Gsk3b* KO and control littermates or in the growth plates of control mice (Fig. 6A, S5A). However, histologic evaluation of the growth plate in KO mice showed presence of TUNEL positive cells exhibiting various features of apoptosis such as cell shrinkage (blue arrows), pyknotic nuclei (black arrows), blebbing of cell membrane (orange arrows), and presence of apoptotic bodies (red arrows) in the tissue. Unlike control mice (Fig. 6A, S5A), matrix staining was also observed in the growth plates of *Gsk3a/Gsk3b* KO mice, which could be due to inefficient clearing of fragmented DNA and dying cell debris during the process of accelerated cell death (Fig. 6A). TUNEL assay was not performed in P43 and P58 mice because of almost complete absence of chondrocytes at these later stages.

**Fig. 6.**
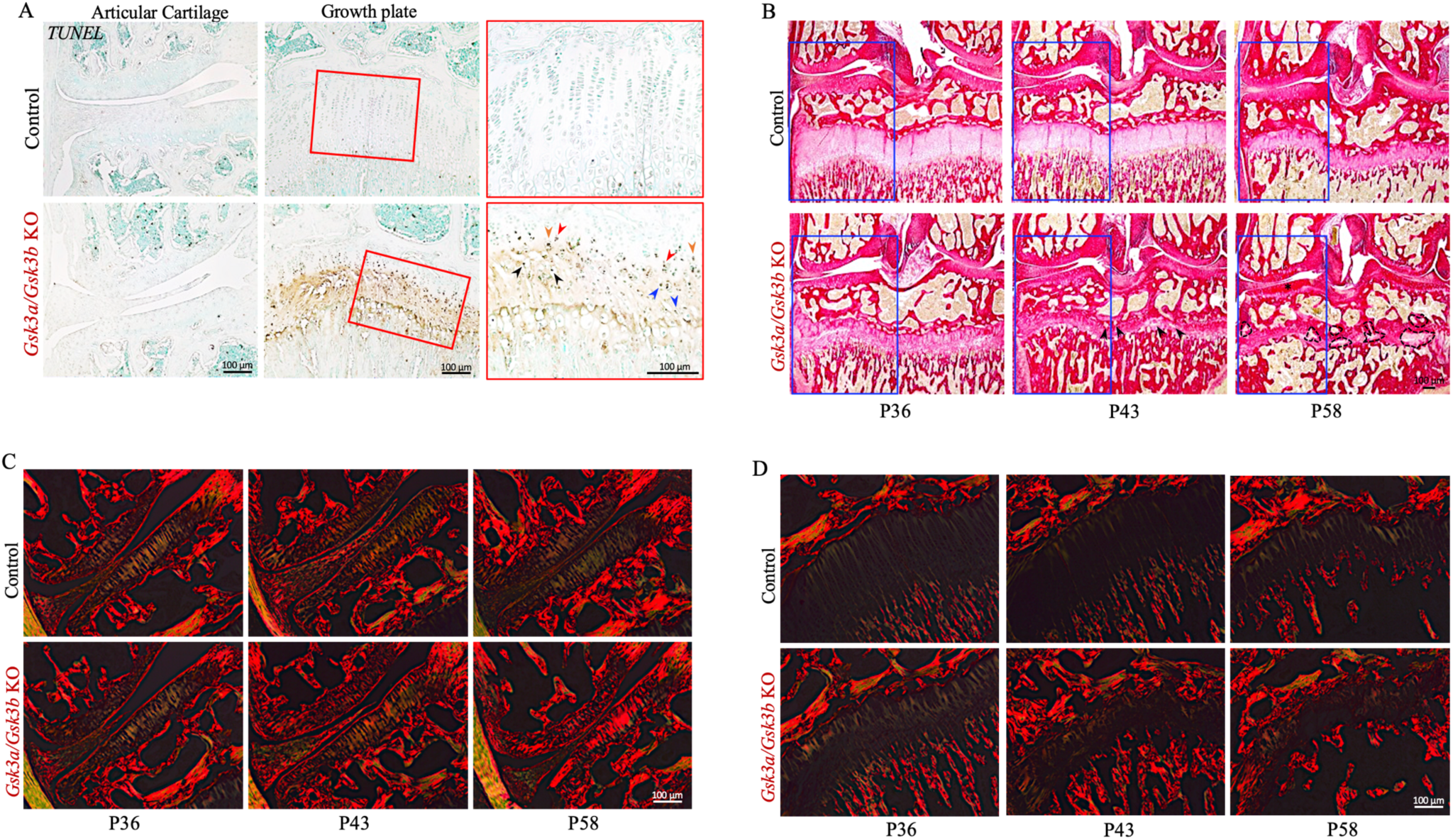
Increased chondrocyte apoptosis and compromised composition of growth plate in postnatal cartilage-specific *Gsk3a/Gsk3b* KO mice. Cell death by apoptosis was detected in P36 *Gsk3a/Gsk3b* KO and control mice by TUNEL assay (A) where various apoptotic cell stages such as shrinking cell, pyknotic nuclei, blebbing of cell membrane and presence of apoptotic bodies are indicated by blue, black, orange and red arrows, respectively. Images are representative of 3-4 mice per group. Picrosirius Red S stained frontal knee joint sections were used to examine the collagen network in *Gsk3a/Gsk3b* KO and control mice at all three time points in brightfield images (B), where bone marrow cavity-like regions are highlighted with black arrows or dotted line, and in linear polarized light images (C&D) of articular cartilage (C) and growth plate (D). Images are representative of 4-7 mice per group.

Since loss of Toluidine Blue stain intensity in KO cartilage indicated proteoglycan loss (Fig. 3), we were curious to further examine the composition of the growth plate. To address this, knee joint sections were stained with Picrosirius Red S to assess collagen content and organization in cartilage. Picrosirius Red S stains collagen in red and cells in yellow colour when imaged in brightfield. In control mice, growth plates showed normal structural organization and stained faintly with Picrosirius Red S in P36 and P43 mice, while staining intensity of growth plates increased at P58, suggesting higher density of collagen fibers in P58 compared to younger control mice (Fig. 6B, S5B). Similarly, *Gsk3a/Gsk3b* KO mice also showed increasing intensity of red colour with the age of growth plate from P36 to P58 (Fig. 6B), and presence of bone marrow-like regions in remnants of growth plate highlighted with black arrows and dotted lines in P43 and P58 KO, respectively. However, at all three time points, growth plates of KO mice stained more intensely with Picrosirius Red S than those of control littermates, suggesting higher density of collagen in KO mice. Similarly, articular cartilage in P58 *Gsk3a/Gsk3b* KO mice stained more intensely with Picrosirius Red S compared to control littermates (Fig. 6B, S5B).

Further analyses of Picrosirius Red S stained sections in linear polarized light displayed a green colour, indicating thinner collagen fibres, in P36 articular cartilage in both *Gsk3a/Gsk3b* KO and control mice compared to later stages, as well as in P43 and P58 control mice compared to KO mice. In P43 KO mice, articular cartilage stained red from superficial to middle zone indicating thicker collagen fibres, followed by green coloured thin collagen fibres in deep zone separating middle zone articular cartilage from subchondral bone. An extended region of intense red colour from superficial to deep zone in articular cartilage, separated from subchondral bone by thin green collagen fibres, was observed in P58 KO mice (Fig. 6C, S5C). The progressively decreasing thickness of growth plates in control mice displayed elongated green to orange coloured thin collagen fibres from P36 to P58. While shrinking growth plates in P36 KO mice revealed the presence of green to orange coloured thin collagen fibres, P43 to P58 growth plates showed progressive disorganization of collagen fibres, increase in orange to red thicker collagen fibres and completely disrupted growth plates in P58 KO mice (Fig. 6D, S5D). These findings support the presence of thicker collagen fibres and its dense network in cartilage of *Gsk3a/Gsk3b* KO mice compared to control littermates.

### Postnatal cartilage-specific *Gsk3a/Gsk3b* KO mice show increased Col2 staining in articular cartilage and growth plate

The above observation of thicker fibrils and dense collagen network in articular cartilage and growth plate remnants of P58 KO mice (Fig. 6 B-D) was exactly opposite to the reduced Toluidine Blue staining for proteoglycans (Fig. 3). To further clarify above findings, immunostaining for Col2 was performed on frontal knee joint sections. Articular cartilage in P36, P43 and P58 control mice stained consistently for the presence of Col2 from superficial to deep zones. Compared to controls, a band of higher intensity staining was observed extending from the superficial to the middle zone of articular cartilage in P36 *Gsk3a/Gsk3b* KO mice, which was further enhanced and extended in P43 and P58 KO mice (Fig. 7A, S6A). Unlike articular cartilage, growth plates in control mice did not stain positive for Col2, possibly since a non-enzymatic detergent-based method of antigen retrieval was used during immunohistochemistry. Growth plates in *Gsk3a/Gsk3b* KO mice showed progressive, positive staining for Col2 from P36 to P58 compared to control mice (Fig. 7B, S6B). Besides increased synthesis or stability of Col2, this increased staining could be due to unmasking of Col2 fibres after proteoglycan loss in KO mice. To validate masking/unmasking effects of proteoglycans on Col2 immunostaining in cartilage, frontal knee joint sections of P58 wildtype mice were treated with chondroitinase to digest glycosaminoglycans, followed by Toluidine Blue staining to detect proteoglycans and immunostaining for Col2. Chondroitinase-treated knee joint sections showed loss of Toluidine Blue staining in articular cartilage and growth plate, confirming loss of proteoglycans. Compared to untreated controls, chondroitinase-treated knee joint sections stained intensely for the presence of Col2 in both articular cartilage and growth plate (Fig. 7C). This staining upon chondroitinase treatment was similar to that of P58 *Gsk3a/Gsk3b* KO cartilage (Fig. 3 A&B, 7 A&B), thus providing evidence for an unmasking effect of proteoglycan loss in cartilage of *Gsk3a/Gsk3b* KO mice.

**Fig. 7.**
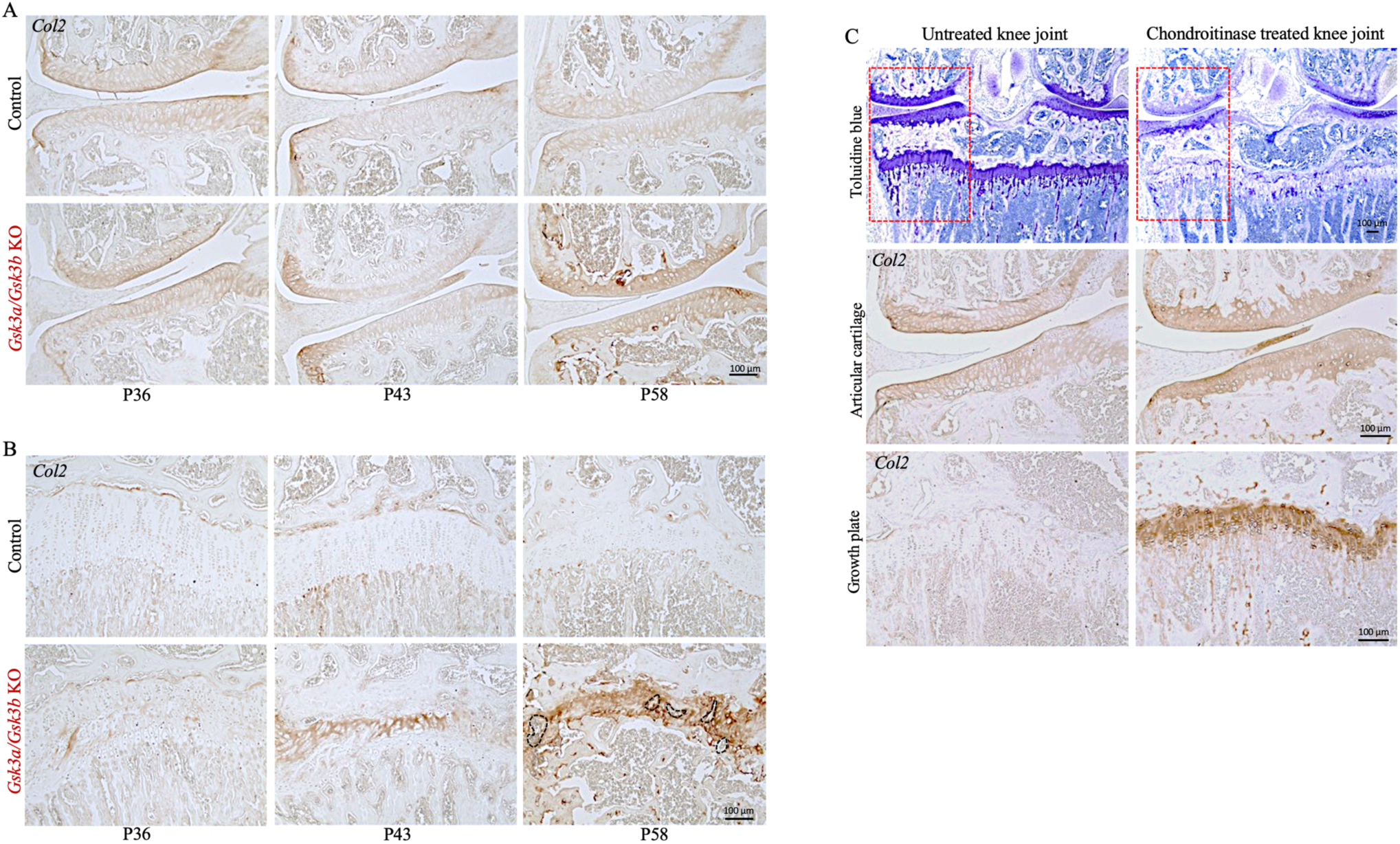
Presence of increased Col2 levels in articular cartilage and growth plate of cartilage-specific *Gsk3a/Gsk3b* KO mice. Representative images of localized Col2 immunostaining in articular cartilage (A) and growth plate (B) of *Gsk3a/Gsk3b* KO and control mice at P36, P43 and P58, assessed by immunohistochemistry using frontal knee joint sections from 4-7 mice for each experimental group. Representative images of chondroitinase-treated or untreated frontal knee joint sections stained with Toluidine Blue and Col2 (C) from four mice per treatment.

### Beta-catenin and GLI1 expression in *Gsk3a/Gsk3b* KO mice

We next examined potential mediators of GSK3 signaling, such as beta-catenin and GLI1. Being downstream of GSK3 and occupying a key node in the canonical Wnt signaling pathway, the expression profile of beta-catenin was investigated in P36 knee joints sections. Basal levels of beta-catenin were expressed in all zones of the growth plate as well as in articular cartilage of control mice (Fig. 8 A&B, S7 A&B). While articular cartilage continued the basal expression of beta-catenin in *Gsk3a/Gsk3b* KO mice, some of the remaining cells scattered at the interface of cartilage-primary spongiosa and in chondrocyte-clusters in the resting zone of the disorganized growth plates stained intensely for beta-catenin in both male (Fig. 8A) and female KO (Fig. 8B) mice.

**Fig. 8.**
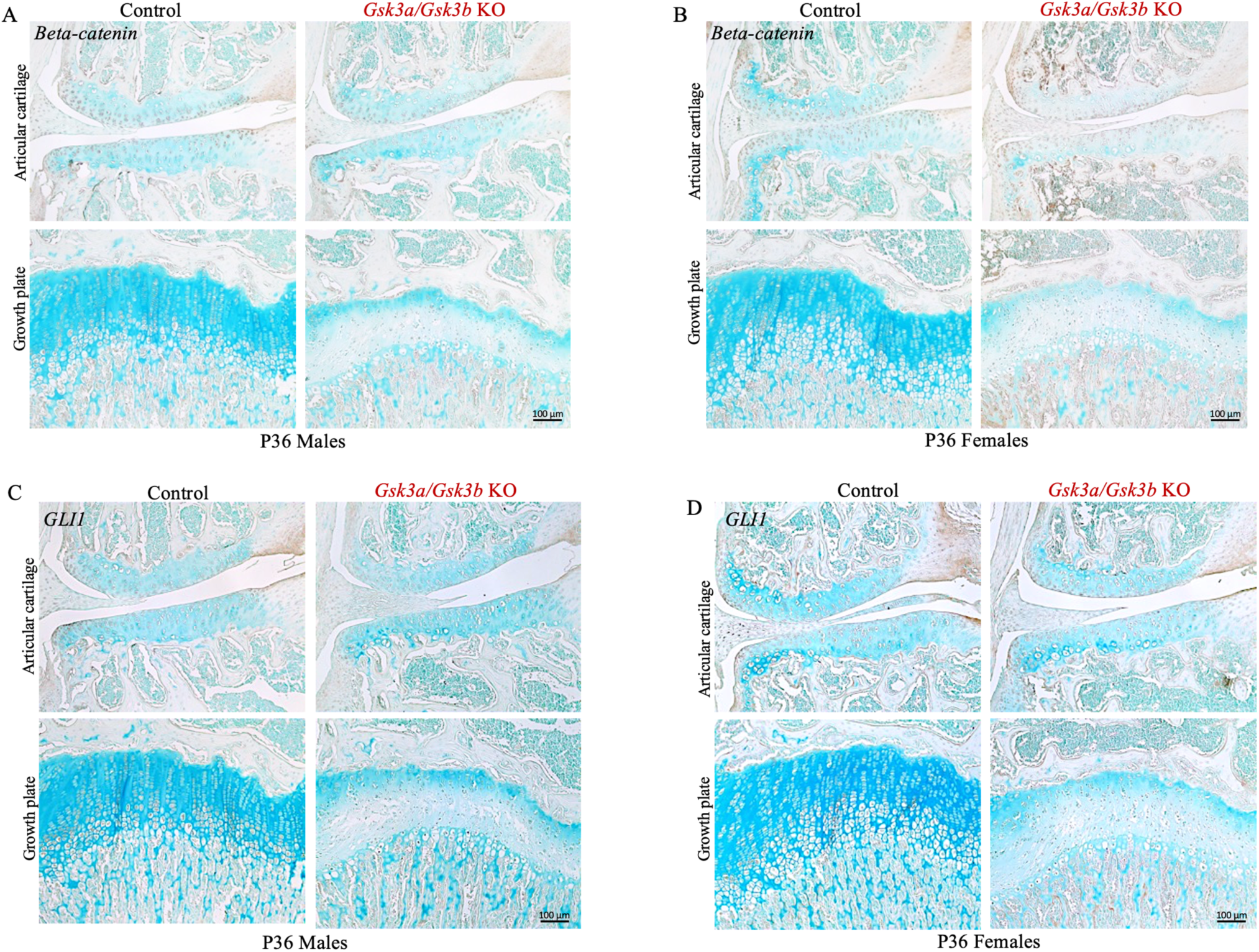
Beta-catenin and GLI1 expression in cartilage-specific *Gsk3a/Gsk3b* KO mice. Localized expression of beta-catenin (A&B) and GLI1 (C&D) was assessed in P36 male (A&C) and female (B&D) *Gsk3a/Gsk3b* KO and control littermates by IHC. Images are representative of 5-7 mice per group.

IHH is another crucial signaling pathway for skeletal development and maintenance, which works in both PTHrP-dependent and -independent ways (25,26). Interaction of these key pathways, along with others, decide the fate of skeletal components and cells. Studies have shown that IHH is crucial to induce columnar chondrocyte differentiation in the growth plate postnatally (14,16,26). To investigate the involvement of IHH signaling in our study model, GLI1- a transcription factor directly regulated by IHH - was examined in knee joint sections of P36 mice. No significant difference in the expression profile of GLI1 was observed in articular cartilage of control and *Gsk3a/Gsk3b* KO mice (Fig. 8 C&D, S7C&D). In controls, GLI1 was expressed in proliferative to pre-hypertrophic chondrocytes in the growth plate. Similar to beta-catenin, the expression of GLI1 was limited to few remaining pre-hypertrophic cells scattered in chondrocyte-clusters in resting zone of disorganized growth plates in both male (Fig. 8C) and female (Fig. 8D) KO mice.

## DISCUSSION

Earlier reports targeting either GSK3alpha or GSK3beta have demonstrated the importance of GSK3 signaling in various conditions of health and disease (3,4,6–8,18,19). During skeletal development, the absence of GSK3beta in cartilage is likely compensated by upregulation of its highly related isoform, GSK3alpha, at least to some extent (6,7). However, conditional targeting of both GSK3 proteins together, has not been studied in this context. Our study, for the first time to our knowledge, genetically targets both GSK3 genes together, in cartilage and shows a crucial role of GSK3 signaling in cartilage biology, its maintenance and growth as well as skeletal development.

Our study shows that postnatal conditional ablation of *Gsk3a/Gsk3b* in mouse cartilage before skeletal maturity compromises their skeletal development and growth, causing growth retardation. During early time points, *Gsk3a/Gsk3b* KO mice grew at a similar rate as control littermates, while a lag in weight gain appeared over time from P36 to P43. A drastic decrease in percent weight gain and smaller skeletal size in addition to lethargy, reduced motility, and inefficient breathing were exhibited by P58 *Gsk3a/Gsk3b* KO mice, which did not survive beyond P59. Analyses of skeletal components demonstrated mineralization of costo-chondral junctions in the ribcage in *Gsk3a/Gsk3b* KO mice. We speculate that this phenotype might be affecting the elasticity of ribcage required for normal breathing, thus resulting in inefficient breathing in KO mice. All the above-mentioned characteristics might be important factors contributing towards failure of survival after cartilage-specific *Gsk3a/Gsk3b* ablation in skeletally immature mice.

Histological analyses of frontal knee joints of our KO mice showed rapid and progressive loss of cells and proteoglycan content, as well as disrupted structural integrity of the growth plate, advancing into precocious remodeling of the growth plate at the latest time point. When knee joint sections were analyzed to characterize the nature of growth plate tissue in *Gsk3a/Gsk3b* KO mice, later time points showed absence of cells, occurrence of osteoclast-mediated resorption and formation of bone marrow cavity-like regions within the remnants of the growth plate, as well as positive staining for the bone marker osteocalcin. However, the extracellular matrix of the growth plate of *Gsk3a/Gsk3b* KO mice retained high levels of Col2, suggesting that cartilage was not completely replaced by bone-like tissue. Picrosirius Red S stained knee joint sections analyzed indicated a higher collagen content in articular cartilage of our KO mice from P36 to P58, while their growth plate remnants showed presence of limited but thicker collagen fibres compared to control mice. When examined for the type of collagen content in cartilage by immunohistochemistry for Col2, increasing staining intensity for Col2 was seen in both articular cartilage and growth plate of *Gsk3a/Gsk3b* KO mice from P36 to P58. One potential explanation for this increased staining in KO mice is that proteoglycans in cartilage of control mice masks collagen, rendering it less available to staining. Loss of proteoglycans exposes collagen, making it available to stain positive with Picrosirius Red S and Col2 antibodies, thereby enhancing stain intensity in KO mice. This unmasking of Col2 network was confirmed by staining performed on chondroitinase-treated knee joint sections of control mice, providing further evidence that increased Col2 staining is due to unmasking rather than increased levels. Alternative explanations for the increased Col2 staining in KO mice are increased synthesis, which appears unlikely given the absence of chondrocytes in these mice, or stability, which also would seem unlikely given the relatively long half-life of Col2 in cartilage of control mice.

Our data suggest that loss of proteoglycans and chondrocytes in conjunction with increased osteoclast activity result in failure to maintain the growth plate in *Gsk3a/Gsk3b* KO mice, despite the continued presence of Col2. Because of the death of KO mice at P59, we could not determine whether there is an eventual degradation of Col2, leading complete resorption of the growth plate, and fusion of primary and secondary ossification centers in KO mice. As death might be due to rib cage deformities, later activation of aggrecan CreER or use of a Cre driver line that is not active in the axial skeleton (e.g. Prx1 CreER) might be possible approaches to tackle this question.

Previous studies on deciphering the role of GSK3 signaling in physiological conditions involved deletion of either *Gsk3a* or *Gsk3b*, which showed high functional redundancy in the regulation of Wnt/beta-catenin signaling (27). Studies on skeletal development involving deletion of *Gsk3b* in cartilage showed upregulation of GSK3alpha, likely compensating for the loss of GSK3beta activity. The same study also reported increased lengths of resting and proliferative zones but a shorter hypertrophic zone in embryonic tibiae upon pharmacological inhibition of both GSK3 proteins ex-vivo (7). Here, we observed a far more severe phenotype in the growth plate leading to its precocious remodeling in *Gsk3a/Gsk3b* KO mice. These results confirm redundancy of the two GSK3 proteins, as these phenotypes are much more dramatic than those of cartilage and bone-specific *Gsk3b* KO mice and global *Gsk3a* KO mice (5–7,18,19). In contrast to growth plate, articular cartilage in *Gsk3a/Gsk3b* KO mice showed much milder features of loss of proteoglycans and Sox9 positive cells, but no structural degeneration. These differential effects of *Gsk3a/Gsk3b* KO in articular cartilage and growth plate cartilage suggests intrinsic differences in both types of cartilage and would be interesting to investigate.

The role of GSK3 as a negative regulator of Wnt/beta-catenin signaling is well established, where shifts in binding of GSK3 to LRP5/6 away from beta-catenin relieves this latter molecule of targeting for ubiquitylation and accumulation of beta-catenin in cells and, thence, activation of downstream genes. Beta-catenin mediates increased expression of matrix metalloproteinases and aggrecanases, which also promotes invasion of endochondral ossification into the growth plate. Increased beta-catenin signaling induces mitochondrial death, accelerated apoptosis in growth plate cells, and proteoglycan loss in both growth plate and articular cartilage, thus deranging the growth plate structure and composition (13,28–30). Previous study has shown that transient activation of Wnt/beta-catenin signaling induces abnormal closure of the growth plate in postnatal mice and thickening of their articular cartilage (13). Our study in *Gsk3a/Gsk3b* KO mice also shows a similar growth plate phenotype, while the articular cartilage phenotype is not recapitulated. This suggests that the growth plate phenotype in our KO mice is mediated by activated canonical Wnt signaling (at least in part), while this plays less of a role in articular cartilage under our experimental conditions. We expected to see increased levels of beta-catenin in our KO mice, which could at least partially explain the observed growth plate phenotype. While we did not find clear evidence for such an effect in the growth plate, this is most likely due to the time points chosen – a time point closer to initial tamoxifen administration might have demonstrated increased beta-catenin signaling. However, no increased staining for beta-catenin was seen in KO articular cartilage despite the clear presence of chondrocytes.

Contrary to beta-catenin (13), deletion of either IHH (14) or PTHrP (15) results in abnormal closure of the growth plate in postnatal mice. Since *Gsk3a/Gsk3b* KO mice also show accelerated remodeling of growth plate compromising longitudinal growth of skeletal elements, we speculate potential, direct or indirect, interaction of IHH and/or PTHrP with GSK3 in cartilage. We chose to examine the IHH-dependent target molecule GLI1, which showed a similar expression profile as beta-catenin in cartilage of both male and female P36 control and KO littermates. In this study, our time points might have been too late to determine the expression of GLI1 in the cartilage of *Gsk3a/Gsk3b* KO mice, similar to beta-catenin.

In summary, the data presented here demonstrate a crucial role of GSK3 signaling in postnatal maintenance of the growth plate. This study opens up multiple avenues to study various direct and indirect targets of GSK3 to enhance basic understanding of cartilage and skeletal tissues and exploit them for translational therapeutics to treat developmental and/or acquired skeletal disorders.

## ACKNOWLEDGEMENTS

We thank all members of our lab for valuable discussions and encouragement, in particular Julia Bowering for tissue sectioning.

## AUTHOR CONTRIBUTIONS

Conceptualization and methodology, S.K.B. and F.B.; investigation, S.K.B., D.B., C.P.; analysis, S.K.B., C.P. and F.B.; manuscript preparation and review, S.K.B., D.B., J.W., and F.B. All authors approved submission of manuscript.

## DECLARATION

The authors declare no conflict of interest.

## REFERENCES

1. DiGirolamo DJ, Clemens TL, Kousteni S. The skeleton as an endocrine organ. Nature Reviews Rheumatology. 2012.

2. Mackie EJ, Ahmed YA, Tatarczuch L, Chen KS, Mirams M. Endochondral ossification: How cartilage is converted into bone in the developing skeleton. International Journal of Biochemistry and Cell Biology. 2008.

3. Beurel E, Grieco SF, Jope RS. Glycogen synthase kinase-3 (GSK3): Regulation, actions, and diseases. Pharmacology and Therapeutics. 2015.

4. Lal H, Ahmad F, Woodgett J, Force T. The GSK-3 family as therapeutic target for myocardial diseases. Circulation Research. 2015.

5. Kaidanovich-Beilin O, Lipina T V., Takao K, Van Eede M, Hattori S, Laliberté C, et al. Abnormalities in brain structure and behavior in GSK-3alpha mutant mice. Mol Brain. 2009;

6. Gillespie JR, Bush JR, Bell GI, Aubrey LA, Dupuis H, Ferron M, et al. GSK-3β function in bone regulates skeletal development, whole-body metabolism, and male life span. Endocrinology. 2013;

7. Gillespie JR, Ulici V, Dupuis H, Higgs A, Dimattia A, Patel S, et al. Deletion of glycogen synthase kinase-3β in cartilage results in up-regulation of glycogen synthase kinase-3α protein expression. Endocrinology. 2011;

8. Cuzzocrea S, Mazzon E, Di Paola R, Muià C, Crisafulli C, Dugo L, et al. Glycogen synthase kinase-3β inhibition attenuates the degree of arthritis caused by type II collagen in the mouse. Clin Immunol. 2006;

9. Yost C, Torres M, Miller JR, Huang E, Kimelman D, Moon RT. The axis-inducing activity, stability, and subcellular distribution of β-catenin is regulated in Xenopus embryos by glycogen synthase kinase 3. Genes Dev. 1996;

10. Kaushal JB, Sankhwar P, Kumari S, Popli P, Shukla V, Hussain MK, et al. The regulation of Hh/Gli1 signaling cascade involves Gsk3β-mediated mechanism in estrogen-derived endometrial hyperplasia. Sci Rep. 2017;

11. Nelson ER, Levi B, Sorkin M, James AW, Liu KJ, Quarto N, et al. Role of GSK-3β in the osteogenic differentiation of palatal mesenchyme. PLoS One. 2011;

12. Sawakami K, Robling AG, Ai M, Pitner ND, Liu D, Warden SJ, et al. The Wnt co-receptor LRP5 is essential for skeletal mechanotransduction but not for the anabolic bone response to parathyroid hormone treatment. J Biol Chem. 2006;

13. Yuasa T, Kondo N, Yasuhara R, Shimono K, Mackem S, Pacifici M, et al. Transient activation of Wnt/β-catenin signaling induces abnormal growth plate closure and articular cartilage thickening in postnatal mice. Am J Pathol. 2009;

14. Maeda Y, Nakamura E, Nguyen MT, Suva LJ, Swain FL, Razzaque MS, et al. Indian Hedgehog produced by postnatal chondrocytes is essential for maintaining a growth plate and trabecular bone. Proc Natl Acad Sci U S A. 2007;

15. Hirai T, Chagin AS, Kobayashi T, Mackem S, Kronenberg HM. Parathyroid hormone/parathyroid hormone-related protein receptor signaling is required for maintenance of the growth plate in postnatal life. Proc Natl Acad Sci U S A. 2011;

16. Maeda Y, Schipani E, Densmore MJ, Lanske B. Partial rescue of postnatal growth plate abnormalities in Ihh mutants by expression of a constitutively active PTH/PTHrP receptor. Bone. 2010;

17. Hoeflich KP, Luo J, Rubie EA, Tsao MS, Jin O, Woodgett JR. Requirement for glycogen synthase kinase-3β in cell survival and NF-κB activation. Nature. 2000;

18. MacAulay K, Doble BW, Patel S, Hansotia T, Sinclair EM, Drucker DJ, et al. Glycogen Synthase Kinase 3α-Specific Regulation of Murine Hepatic Glycogen Metabolism. Cell Metab. 2007;

19. Patel S, Doble BW, MacAulay K, Sinclair EM, Drucker DJ, Woodgett JR. Tissue-Specific Role of Glycogen Synthase Kinase 3 in Glucose Homeostasis and Insulin Action. Mol Cell Biol. 2008;

20. Wang G, Woods A, Agoston H, Ulici V, Glogauer M, Beier F. Genetic ablation of Rac1 in cartilage results in chondrodysplasia. Dev Biol. 2007;

21. Suzuki D, Yamada A, Amano T, Yasuhara R, Kimura A, Sakahara M, et al. Essential mesenchymal role of small GTPase Rac1 in interdigital programmed cell death during limb development. Dev Biol. 2009;

22. Fang H, Huang L, Welch I, Norley C, Holdsworth DW, Beier F, et al. Early Changes of Articular Cartilage and Subchondral Bone in The DMM Mouse Model of Osteoarthritis. Sci Rep. 2018;

23. Dupuis H, Pest MA, Hadzic E, Vo TX, Hardy DB, Beier F. Exposure to the rxr agonist sr11237 in early life causes disturbed skeletal morphogenesis in a rat model. Int J Mol Sci. 2019;

24. Pest MA, Russell BA, Zhang YW, Jeong JW, Beier F. Disturbed cartilage and joint homeostasis resulting from a loss of mitogen-inducible gene 6 in a mouse model of joint dysfunction. Arthritis Rheumatol. 2014;

25. Kobayashi T, Chung U Il, Schipani E, Starbuck M, Karsenty G, Katagiri T, et al. PTHrP and Indian hedgehog control differentiation of growth plate chondrocytes at multiple steps. Development. 2002;

26. Kobayashi T, Soegiarto DW, Yang Y, Lanske B, Schipani E, McMahon AP, et al. Indian hedgehog stimulates periarticular chondrocyte differentiation to regulate growth plate length independently of PTHrP. J Clin Invest. 2005;

27. Doble BW, Patel S, Wood GA, Kockeritz LK, Woodgett JR. Functional Redundancy of GSK-3α and GSK-3β in Wnt/β-Catenin Signaling Shown by Using an Allelic Series of Embryonic Stem Cell Lines. Dev Cell. 2007;

28. Holzer T, Probst K, Etich J, Auler M, Georgieva VS, Bluhm B, et al. Respiratory chain inactivation links cartilage-mediated growth retardation to mitochondrial diseases. J Cell Biol. 2019;

29. Tamamura Y, Otani T, Kanatani N, Koyama E, Kitagaki J, Komori T, et al. Developmental regulation of Wnt/β-catenin signals is required for growth plate assembly, cartilage integrity, and endochondral ossification. J Biol Chem. 2005;

30. Guidotti S, Minguzzi M, Platano D, Santi S, Trisolino G, Filardo G, et al. Glycogen Synthase Kinase-3β Inhibition Links Mitochondrial Dysfunction, Extracellular Matrix Remodelling and Terminal Differentiation in Chondrocytes. Sci Rep. 2017;

