## Supplementary info for "Glycogen synthase kinase 3 alpha/beta deletion induces precocious growth plate remodeling and cell loss in mice"

### SUPPLEMENTARY INFORMATION

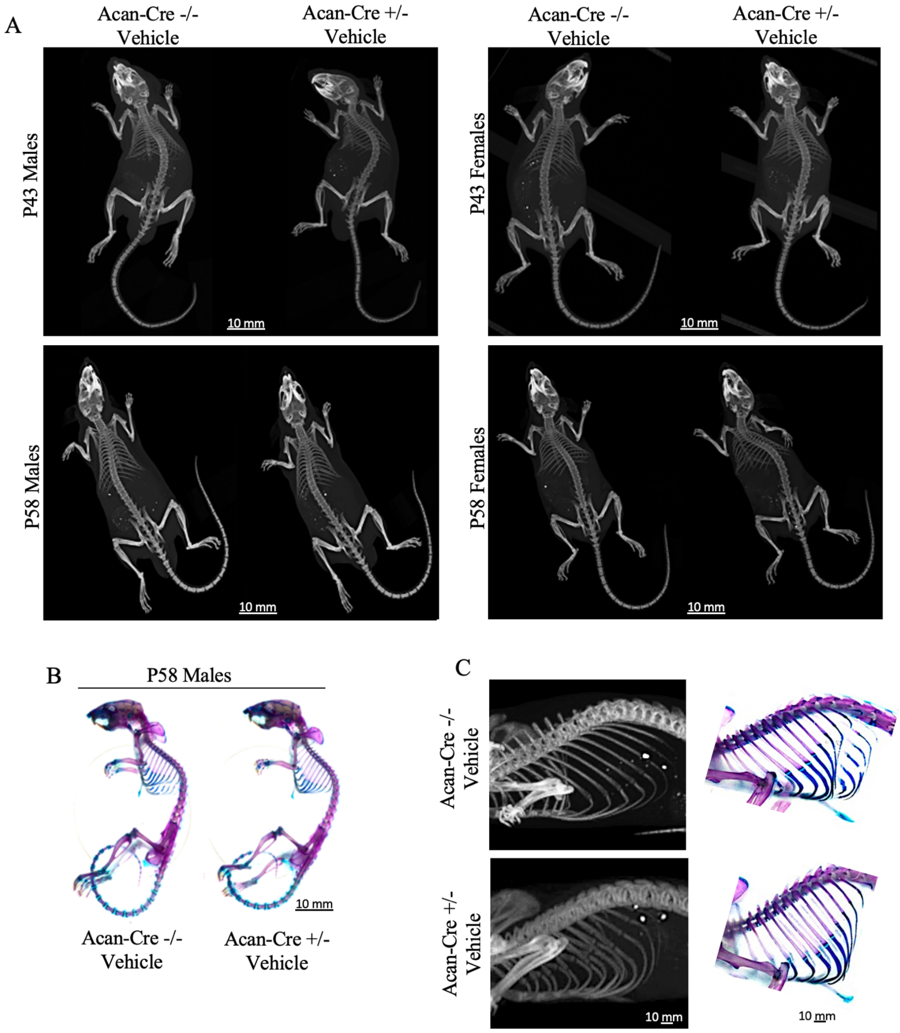


**Suppl. Fig. 1.** Two dimensional micro-CT images of mouse skeletons at P43 and P58 time points in male and female vehicle-treated *Gsk3a*$^{fl/fl}$/*Gsk3b*$^{fl/fl}$ *AcanCre*$^{ERT2+/-}$ and *Gsk3a*$^{fl/fl}$/*Gsk3b*$^{fl/fl}$ littermates (A). Images are representative of 7-13 mice per group. Representative images from six mice per group for Alizarin Red S and Alcian Blue stained mouse skeletons for comparison of bone and cartilage components at P58 in vehicle-treated control littermates (B). Cropped and enlarged region of three dimensional isosurface reconstruction of mouse ribcage highlighting costochondral junctions and supporting skeletal staining in P58 vehicle-treated control littermates (C).

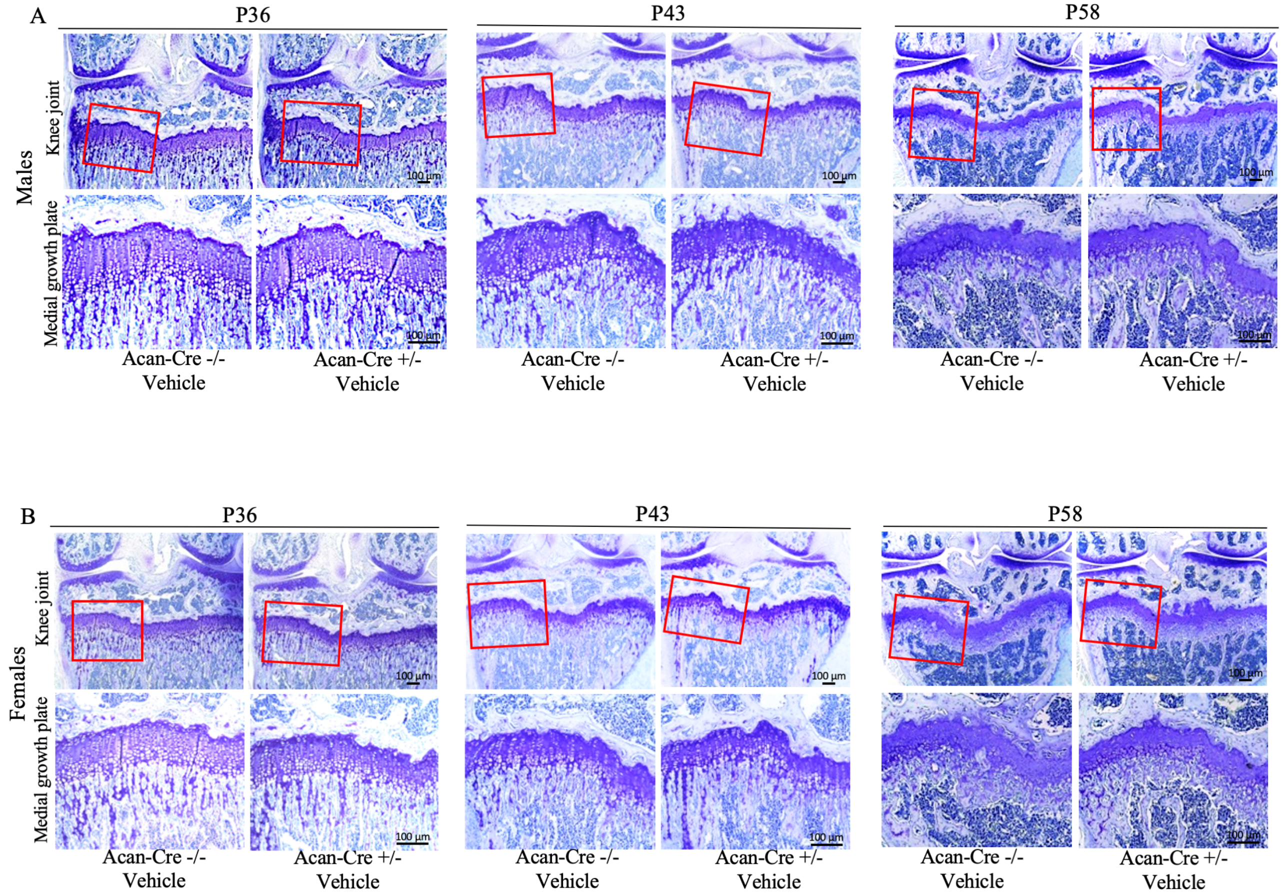


**Suppl. Fig. 2.** Representative images of Toluidine Blue stained frontal knee joint sections from male (A) and female (B) vehicle-treated *Gsk3a*$^{fl/fl}$/*Gsk3b*$^{fl/fl}$ *AcanCre*$^{ERT2+/-}$ and *Gsk3a*$^{fl/fl}$/*Gsk3b*$^{fl/fl}$ littermates at P36, P43 and P58 show no difference compared to tamoxifen-treated Cre negative mice of the same genotype. Images are representative of 4-7 mice per group.

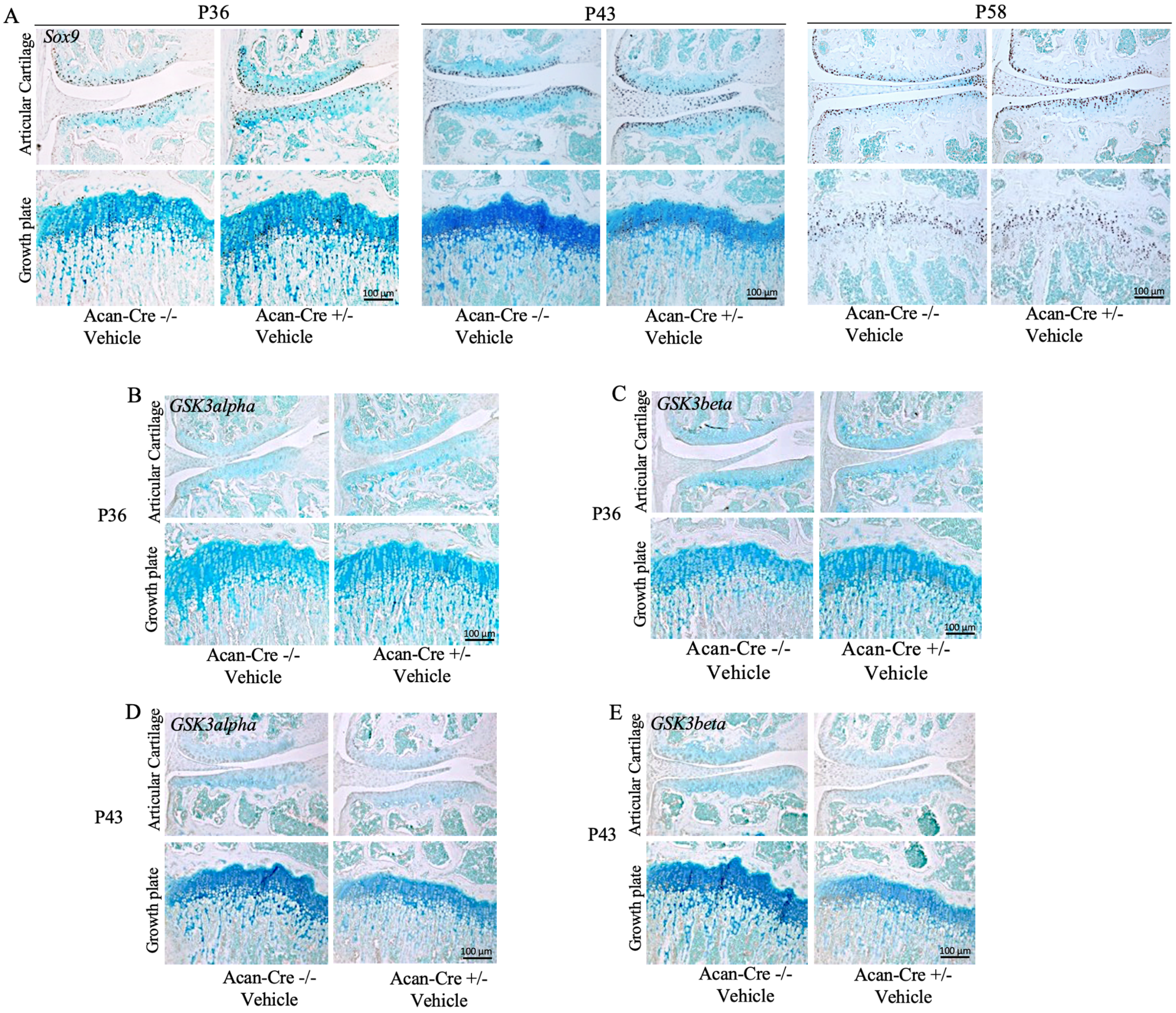


**Suppl. Fig. 3.** Immunohistochemistry on frontal knee joint sections of vehicle-treated $Gsk3a^{fl/fl}/Gsk3b^{fl/fl}$ $AcanCre^{ERT2+/-}$ and $Gsk3a^{fl/fl}/Gsk3b^{fl/fl}$ mice for Sox9 at all three time points (A), and GSK3alpha (B&D) and GSK3beta (C&E) at P36 (B&C) and P43 (D&E) respectively, shows no difference compared to tamoxifen-treated Cre negative mice of the same genotype. Images are representative of 4-7 mice per group for Sox9, and GSK3alpha and GSK3beta represents observations from three mice per group, respectively.

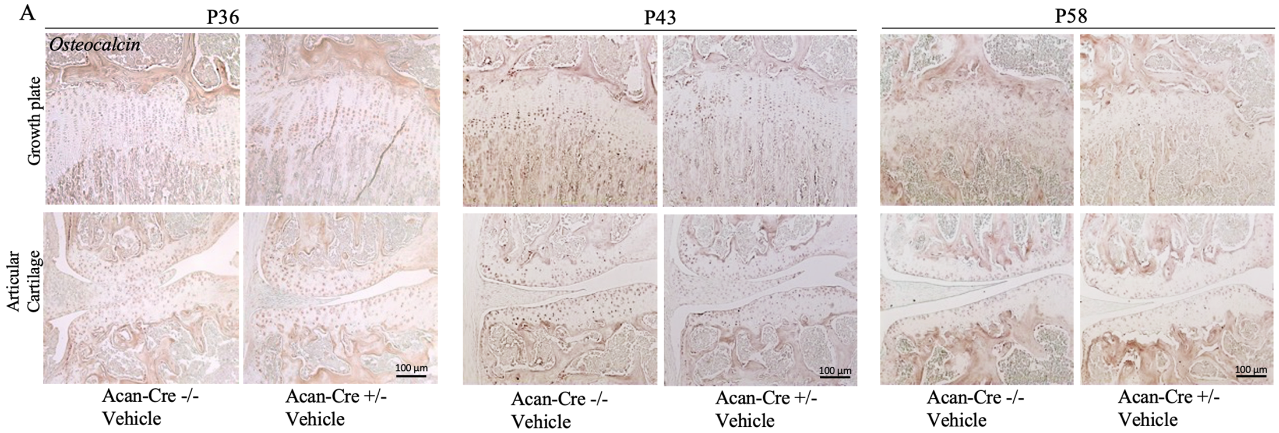


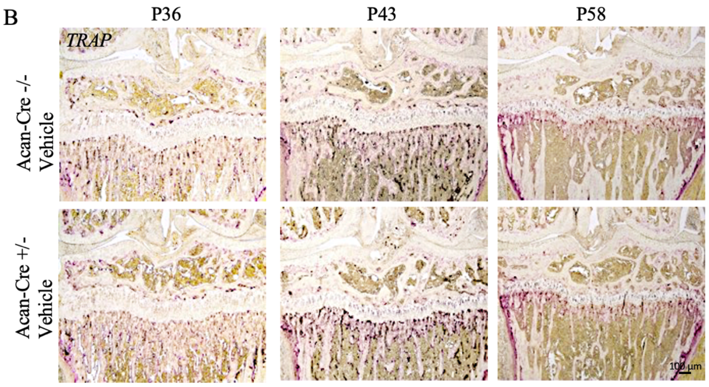


**Suppl. Fig. 4.** Vehicle-treated *Gsk3a$^{fl/fl}$/Gsk3b$^{fl/fl}$ AcanCre$^{ERT2+/-}$* and *Gsk3a$^{fl/fl}$/Gsk3b$^{fl/fl}$* littermates were assessed for the presence of osteocalcin at P36, P43 and P58 (A). Active growth plate remodeling was determined by staining for TRAP positive osteoclasts at all three time points, which showed basal level of osteoclast activity (B). Images are representative of 4-7 mice per group for osteocalcin, and 3-4 mice per group for TRAP staining.

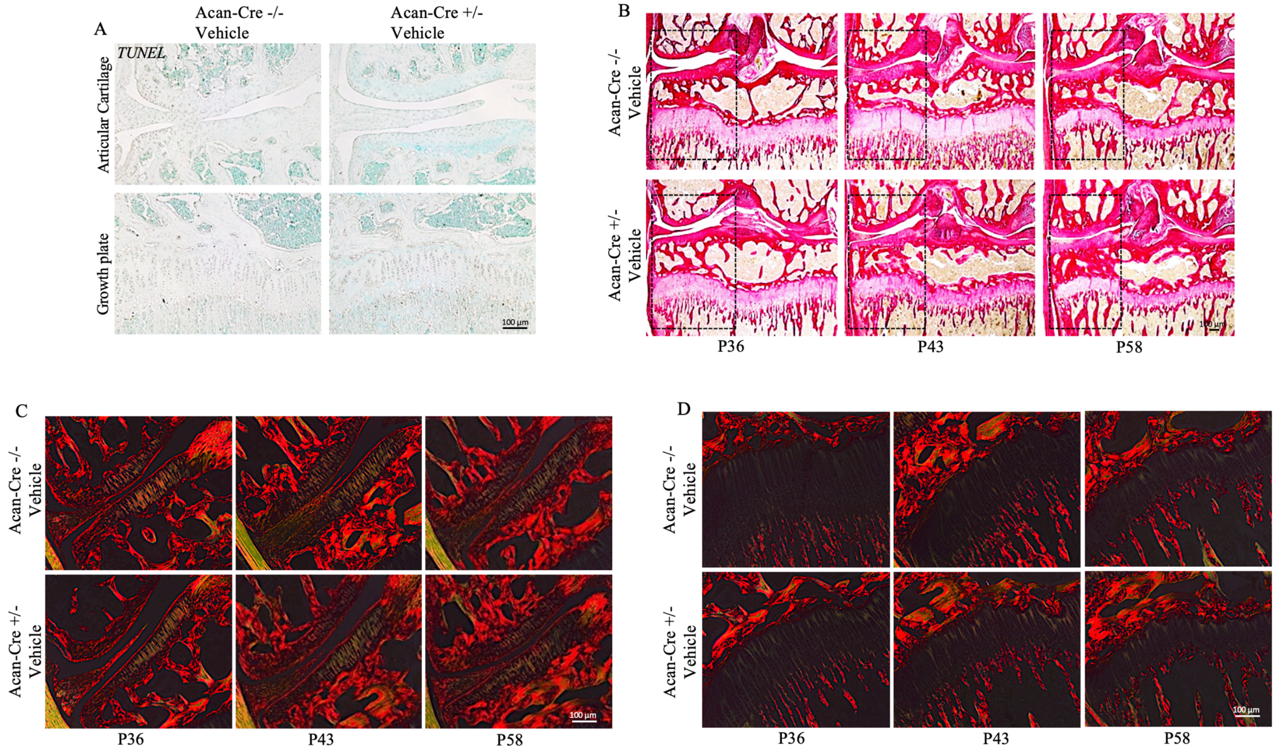


**Suppl. Fig. 5.** TUNEL assay performed on vehicle-treated $Gsk3a^{fl/fl}/Gsk3b^{fl/fl}$ $AcanCre^{ERT2+/-}$ and $Gsk3a^{fl/fl}/Gsk3b^{fl/fl}$ littermates at P36 (A) showed absence of apoptotic cells in articular cartilage and growth plate of these control mice. Images are representative of three mice per group. Frontal knee joint sections showed normal healthy tissue and collagen network in brightfield images (B) as well as in linear polarized light images (C&D) of articular cartilage (C) and growth plate (D) upon Picrosirius Red S staining. Images are representative of 4-7 mice per group.

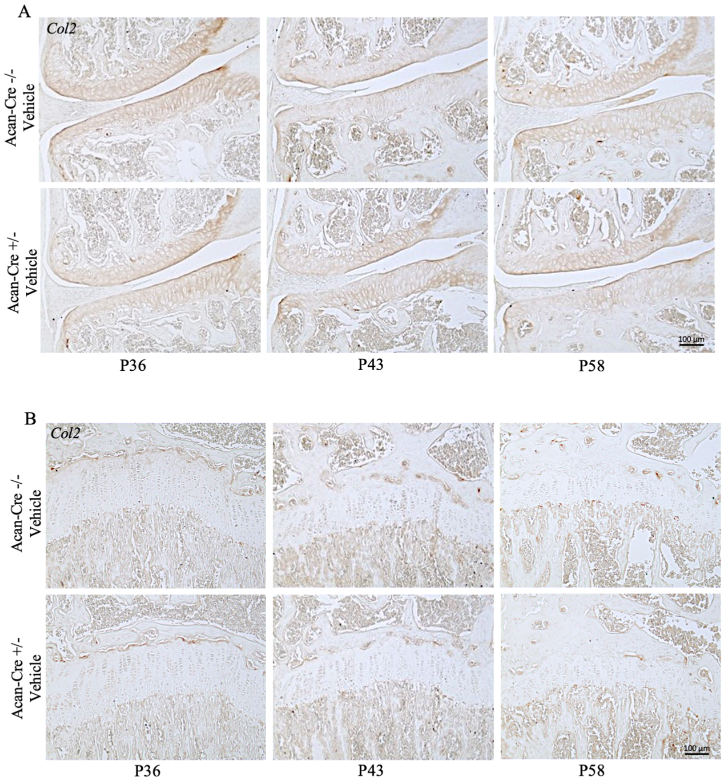


**Suppl. Fig. 6.** Vehicle-treated *Gsk3a*$^{fl/fl}$/*Gsk3b*$^{fl/fl}$ *AcanCre*$^{ERT2+/-}$ and *Gsk3a*$^{fl/fl}$/*Gsk3b*$^{fl/fl}$ littermates were assessed for the presence of Col2 in articular cartilage (A) and growth plate (B) at P36, P43 and P58 by immunohistochemistry using frontal knee joint sections from 4-7 mice for each experimental group.

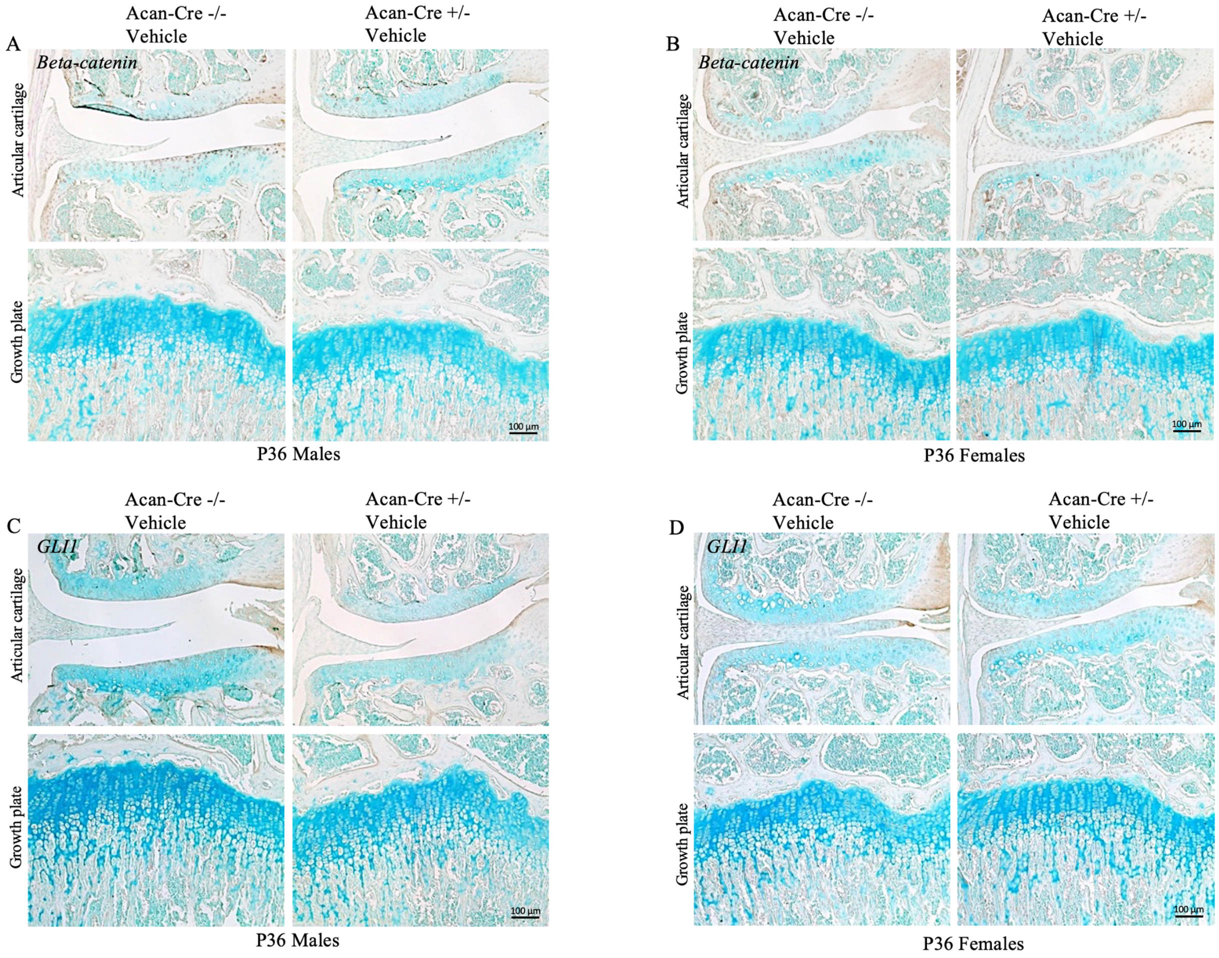


**Suppl. Fig. 7.** IHC for beta-catenin (A&B) and GLI1 (C&D) in P36 vehicle-treated male (A&C) and female (B&D) *Gsk3a*$^{fl/fl}$/*Gsk3b*$^{fl/fl}$ *AcanCre*$^{ERT2+/-}$ and *Gsk3a*$^{fl/fl}$/*Gsk3b*$^{fl/fl}$ littermates showed similar expression pattern as tamoxifen-treated Cre negative mice of the same genotype, as explained in figure 8 A-D, respectively. Images are representative of 5-7 mice per group for both beta-catenin and GLI1, respectively.
